# Genetic and genomic analysis of oxygen consumption in mice

**DOI:** 10.1101/2022.02.05.479269

**Authors:** Shinichiro Ogawa, Hongyu Darhan, Keiichi Suzuki

## Abstract

We estimated genetic parameters of oxygen consumption (OC), OC per metabolic body weight (OCMBW), and body weight at three through eight weeks of age in divergently selected mice populations, with an animal model considering maternal genetic, common litter environmental, and cytoplasmic inheritance effects. Cytoplasmic inheritance was considered based on maternal lineage information. For OC, estimated direct heritability was moderate (0.32) and estimated maternal heritability and proportion of the variance of cytoplasmic inheritance effects to the phenotypic variance were very low (both <0.03), implying that causal genes for OC could be located on autosomes. To assess this hypothesis, we attempted to identify possible candidate causal genes by performing pool-seq using pooled DNA samples from mice in high and low OC lines and selective signature detection. We made a list of possible candidate causal genes for OC, including those relating to electron transport chain and ATP-binging proteins (*Ndufa12, Sdhc, Atp10b*, etc.), *Prr16* encoding Largen protein, *Cry1* encoding a key component of the circadian core oscillator, and so on. The results could contribute to elucidate the genetic mechanism of OC, an indicator for maintenance energy requirement and therefore feed efficiency.

## 1 INTRODUCTION

Reducing feeding costs, occupying more than half of livestock production costs, becomes more important for sustainable development in livestock industry. As an indicator for feeding cost, increasing attention has been focused on residual feed intake (RFI; Koch et al., 1963) (e.g., Gilbert et al., 2017; Hill & Azain, 2009; Hoque & Suzuki, 2009; Nkrumah et al., 2007). RFI has been estimated to be heritable in several livestock species (e.g., Aggrey et al., 2010; Do et al., 2013; Homma et al., 2021; Okanishi et al., 2008; Takeda et al., 2018). Previous studies have challenged to elucidate the biological mechanism of RFI (e.g., Gondret et al., 2017; Okada et al., 2018; Takeda et al., 2020; Weber et al., 2016), some reporting the importance of mitochondrial function (e.g., Fonseca et al., 2015; Li et al., 2021; Yang et al., 2021).

Factors affecting the individual difference in RFI include maintenance energy requirement (MER) (Herd & Arthur, 2009), and the proportion of MER in total energy intake has been reported to be about 70% to 75% in beef cattle (Ferrell & Jenkins, 1985), and 66% and 78.7% in growing and mature pigs, respectively (Zhang et al., 2014). Hotovy et al. (1991) estimated the heritability of MER to be moderate in cattle. Herd and Bishop (2000) estimated the genetic correlations of RFI to be high positive with MER-related traits in British Hereford cattle population. Therefore, genetic improvement for MER seems to be possible and is expected to contribute to efficient livestock production. On the other hand, directly measuring MER is difficult, thus indirect selection using an indicator trait may be more practical. Nielsen et al. (1997a,b) established two mice lines differing in MER by the divergent selection for heat loss, as an indicator for MER, in mature mice. Line differences have been observed in several kinds of phenotypes, including feed and mitochondrial efficiencies (Kgwatalala & Nielsen, 2004; McDonald & Nielsen, 2007; McDonald et al., 2009). However, measuring heat loss using calorimeter is not timesaving (Nielsen et al., 1997b).

Since MER can be indirectly estimated using oxygen consumption (OC) (Kleiber, 1975), an apparatus has been created to measure OC of mice quickly (Hong et al., 2013,2015) and then the divergent selection experiment was started to create higher and lower OC mice lines (HOC and LOC lines, respectively) (Figures 1, 2). Hong et al. (2013) reported that heritability estimated using the data obtained from the first to ninth generations (G1 to G9) was moderate to high and that there was a significant difference in feed conversion ratio between lines at G8 and G9. Using mice at G16, Darhan et al. (2017) reported that MER estimated based on the method of Eggert and Nielsen (2006) tended to be higher in HOC line, that fat percentage was significantly higher in LOC lime, and that moisture, ash, and protein percentage were significantly higher in HOC line. Using mice at several generations before G17, Hong et al., (2015) and Darhan et al. (2019) showed that feed intake, feed conversion rate, and RFI were significantly lower in LOC line mice, together with revealing the difference in mitochondrial respiration activity between the lines. Nonetheless, it remains unresolved whether the observed change in mitochondrial respiration activity could be explained by the inheritance of autosome or mitochondrial DNA (or perhaps both or none).

**FIGURE 1.**
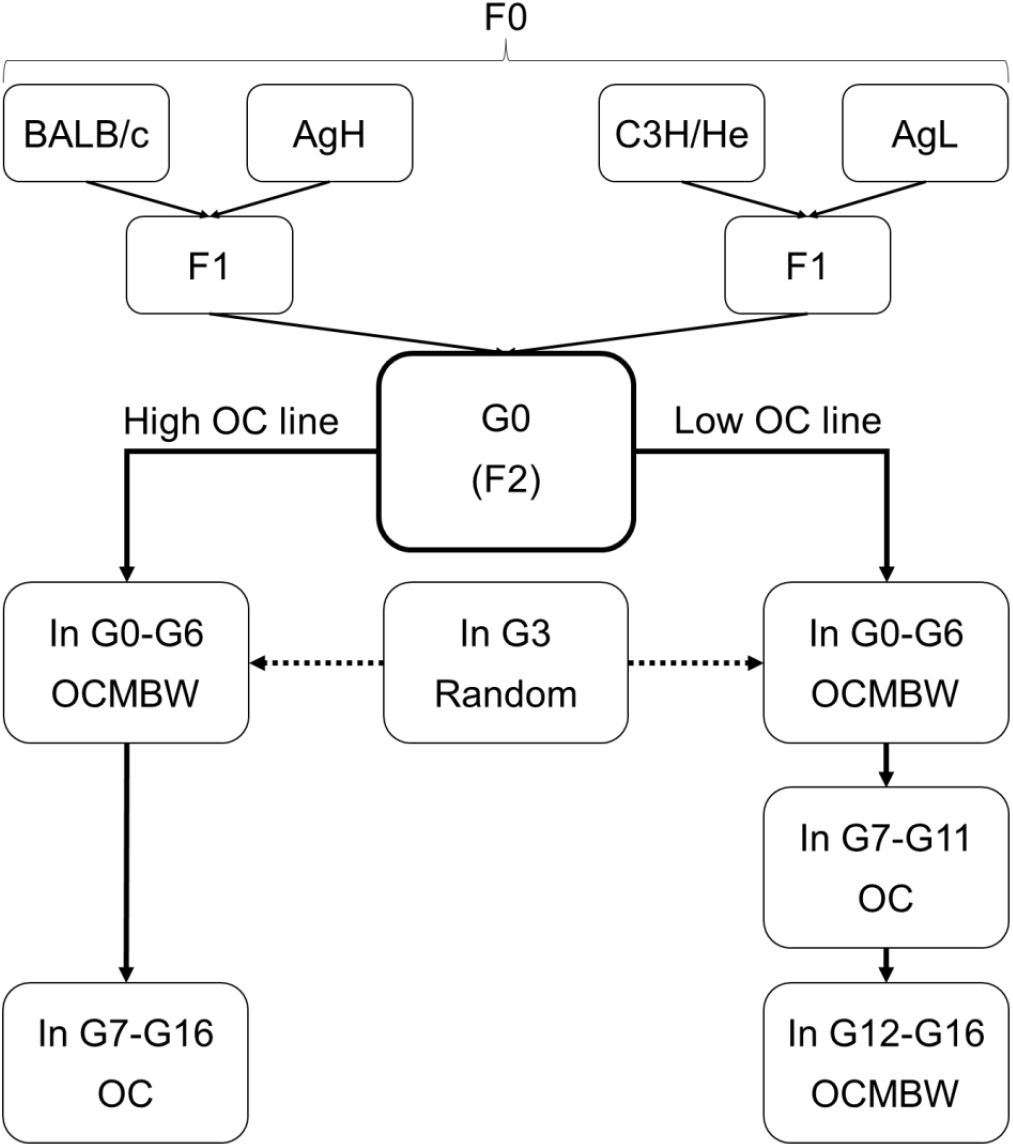
Overviews of schemes of selection experiments for high and low oxygen consumption (OC) performed in previous studies (Darhan et al., 2017, 2019; Hong et al., 2013, 2015). OCMBW: OC per metabolic body weight; AgH and AgL: high and low aggressive lines established by divergent selection by Kawamoto et al. (1993).

**FIGURE 2.**
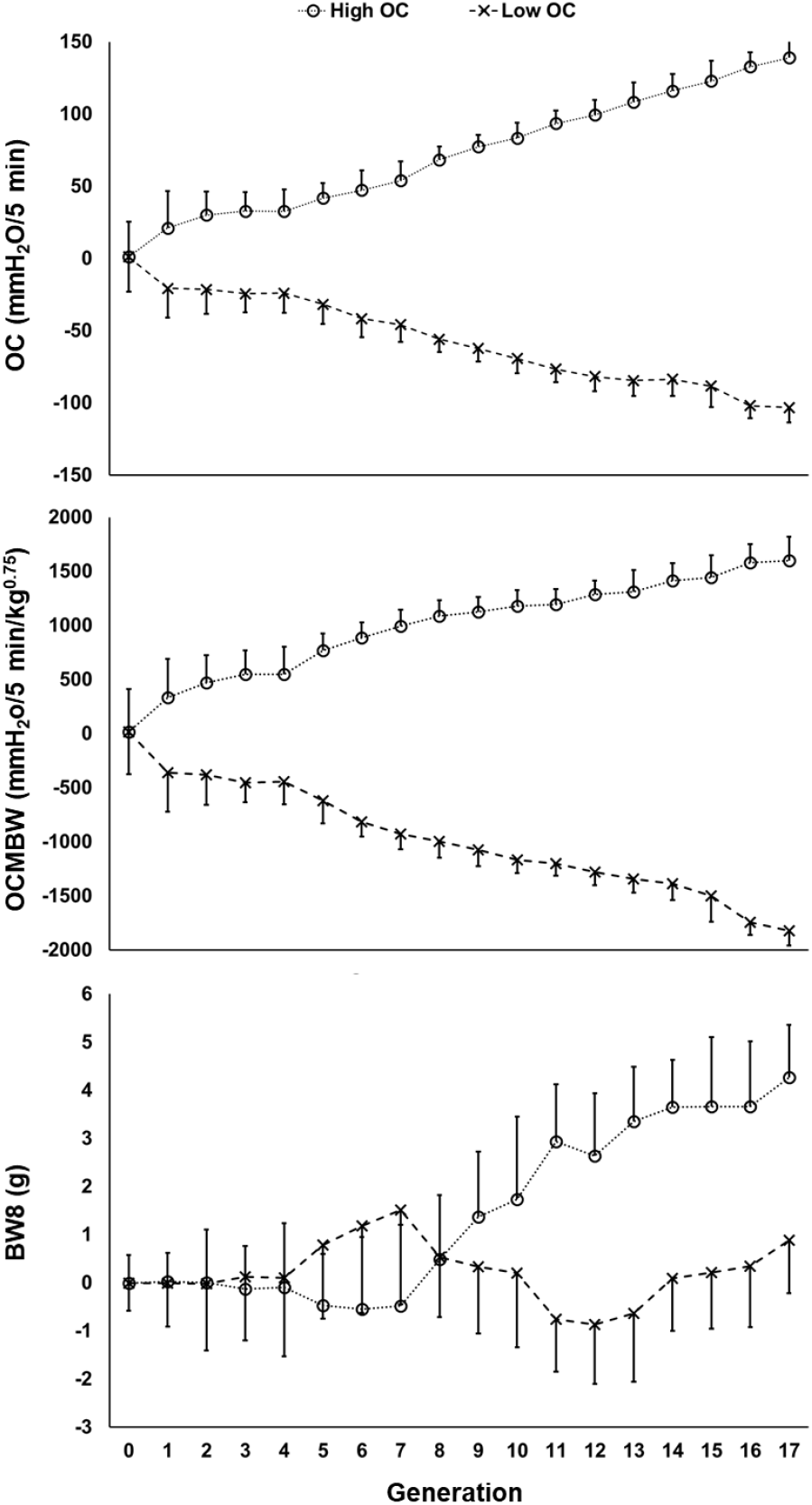
Values of averages and standard deviations (error bars) of estimated breeding values for oxygen consumption (OC), OC per metabolic body weight (OCMBW), and body weight at 8 weeks of age (BW8) per line per generation.

Association between MER and mitochondrial activity is high, approximately 90% of OC is from mitochondria (Rolfe & Brown, 1997). Mitochondrial DNA encodes a part of subunits of respiratory chain complexes and is thought to be maternally inherited, unlike nuclear genome. Therefore, it may be necessary to consider the effect of mitochondrial inheritance in genetic evaluation for OC. On the other hand, there are reports estimating the genetic parameters of body weight in mice with an animal model considering maternal effects (Beniwal et al., 1992; formoso-Rafferty et al., 2016; Leamy et al., 2005). Since body weight affects MER, there may be the impact of maternal effects in OC in mice. However, Hong et al. (2013) did not consider maternal effects in genetic parameter estimation. Identifying novel factors affecting OC is expected to contribute to elucidate the biological mechanism of OC and to set more proper statistical model in genetic evaluation for OC.

Results of selective signature detection might provide important information on the mechanism of phenotype expression (e.g., Andersson, 2012; Nielsen, 2005; Vitti et al., 2013). For example, Castro et al. (2019) performed selective sweep analysis to a mouse population selected over 20 generations for longer tibiae relative to body mass and uncovered many genes, possibly thousands, all acting in concert to increase tibia length. Recently, selective signature analyses have been conducted for various species by using pool-seq data (e.g., Boitard et al., 2012; Kofler et al., 2012; Rubin et al., 2010). Pool-seq is expected to be cost-effective for screening compared to individual sequencing (Anand et al., 2016). We found cryopreserved tail samples of some mice in HOC and LOC lines, which might be useful resources for elucidating the genetic background of OC by pool-seq and selective sweep analysis as a first step.

In this study, we first estimated genetic parameters of OC and related traits with animal models ignoring and considering maternal genetic, common litter environmental, and cytoplasmic inheritance effects. Second, we tried to explore the possible candidate causal genes for OC by pool-seq and selective signature detection analysis.

## 2 MATERIALS AND METHODS

### 2.1 Ethics statement

Selection experiment was performed during the 2010 to 2014 according to Regulations for Animal Experiments and Related Activities at Tohoku University. All mice were reared and handled according to protocol approved by the Institutional Animal Care and Use Committee of Tohoku University.

### 2.2 Selection experiment for oxygen consumption

Detailed explanation is in Hong et al. (2015). Briefly, the base population, G0, consisted of 227 mice, which were F2 population of four-way crossing using BALB/c, C3H, and high and low aggressive lines established by divergent selection (Kawamoto et al., 1993) (Figure 1). Selection objective was to establish HOC and LOC lines while unchanging body weight (BW) as much as possible. Phenotypic records for OC were obtained after fasting all day with water provided *ad libitum*. G1 of each line was obtained by mating 25 males and 25 females selected based on phenotypic records for OC per metabolic BW (= BW^0.75^) at measuring OC (OCMBW). Indicators for truncation selection were estimated breeding values (EBV) for OCMBW from G1 to G6 and EBV for OC in G7 and later in HOC line, and EBV for OCMBW from G1 to G6 and from G12 to G16 and EBV for OC from G7 to G11 in LOC line (Figure 1). Indicator trait was not constant, being apprehensive of correlated response in BW (Hong et al., 2013). EBV were calculated per lines using an animal model considering the effects of sex, generation, and age at measuring OC (linear) and VCE6 program (Neumaier & Groeneveld, 1998). In each generation, OC of all selection candidates (about 150 mice in each line) was measured, and 25 males and 25 females were selected in each line. 25 mating pairs was determined to minimize the inbreeding coefficients of progenies as much as possible. Inbreeding coefficients were calculated using CoeFR program (Satoh, 2000). Age at mating was 12 weeks in average, and only progenies from first parturition were used selection candidates. The number of pups was adjusted to six (three pups in each sex) within a day from parturition until G1 and at seven days of age of pups after G2 when number born alive was larger than six. Cross-fostering was not implemented. All pups were weaned at 21 days of age and reared with feed and water provided *ad libitum* after weaning.

### 2.3 Phenotype and pedigree data

Phenotypic measurements for OC, OCMBW, and BW measured at three through eight weeks of age (BW3, BW4, BW5, BW6, BW7, and BW8) of 4,670 mice derived from 774 dams were analyzed. Note that this study used only records of OC and OCMBW measured after 8 weeks of age to eliminate the effect of fasting at measuring OC on BW. Average values of phenotypic measurements are listed in Table 1. Figure 3 shows averages and standard deviations of OC, OCMBW, and BW8 per generation in each line. In G3, OC could not be measured due to equipment failure (Hong et al., 2013). Corresponding to the differences in selection indices, differences were observed after G4 in OCMBW and after G7 in OC between two lines. In LOC line, average values of OC and OCMBW did not decrease over generations, and similar results have been observed in selection experiments about heat loss in mice (Nielsen et al., 1997b) and OC in chickens at 3weeks of age (MacLaury & Johnson, 1972). Individuals with lower EBVs for OC and OCMBW could have lower basal metabolic rate (Hong et al., 2015), and might have an increased risk of mortality.

**FIGURE 3.**
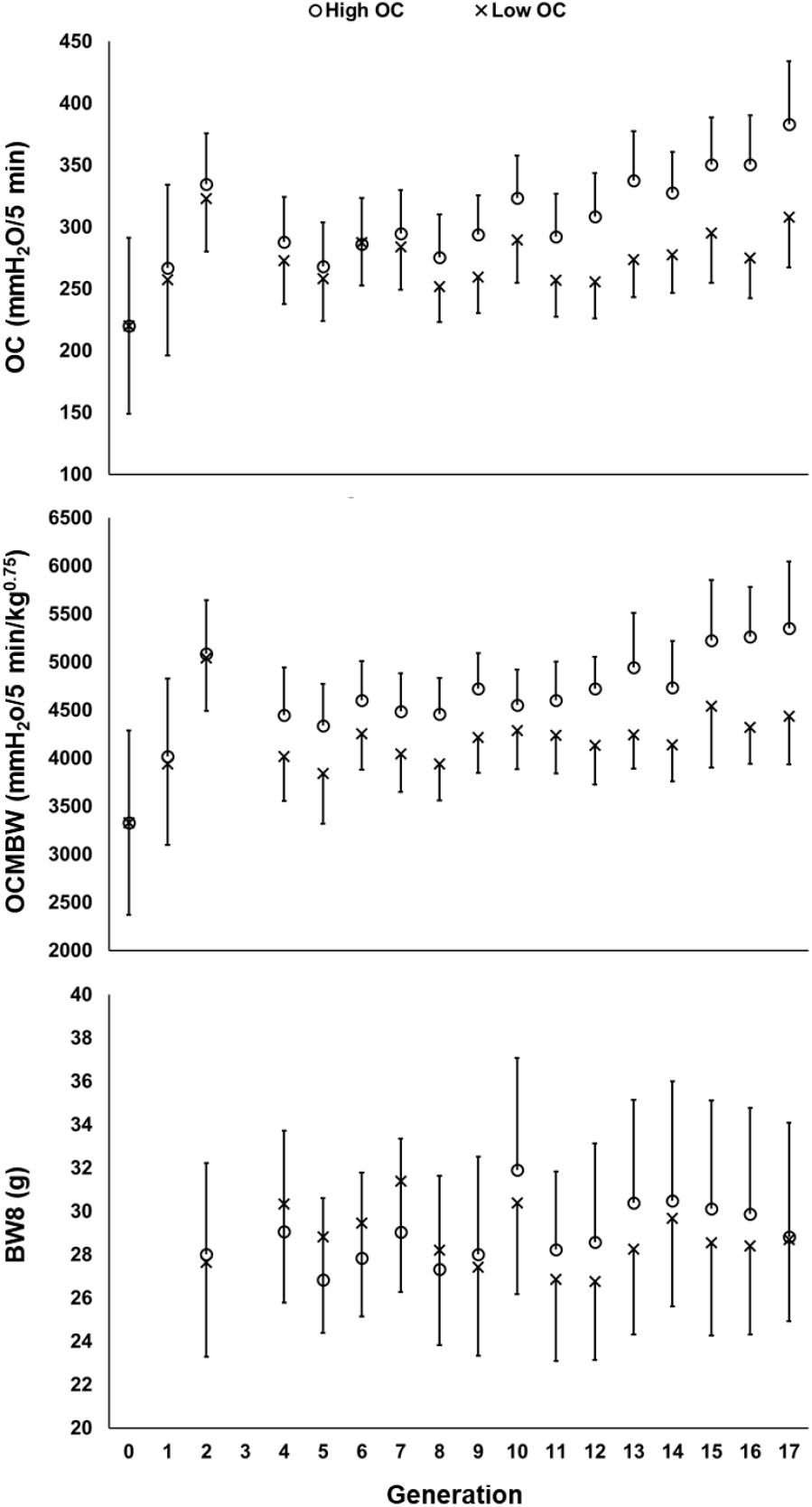
Values of averages and standard deviations (error bar) of phenotypic records for oxygen consumption (OC), OC per metabolic body weight (OCMBW), and body weight at 8 weeks of age (BW8) per line per generation.

**TABLE 1.**
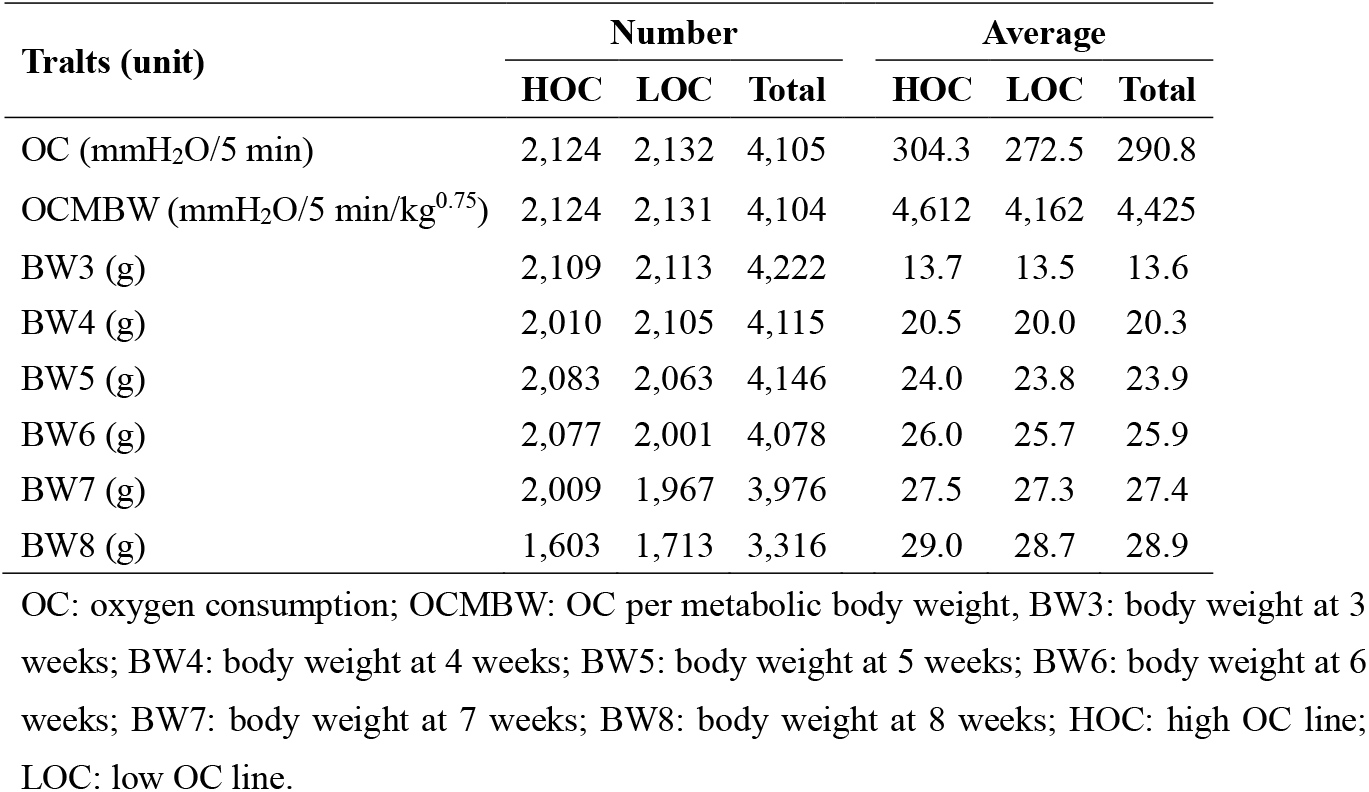
Numbers and average values of phenotypic records of traits studied

Pedigree data for HOC and LOC lines included 2,611 and 6,597 mice, respectively. By merging the two files, a larger pedigree data including 4,878 mice were prepared. Base population of this pedigree data consisted of 21 male and 22 female mice in F0 population (Figure 1). The 22 F0 female mice were used as indicators for maternal inheritance (maternal lineage) in this study. G0 mice with their own phenotypic records belonged to one of the 17 maternal lineages. On the other hand, G17 mice with their own phenotypic records belonged to only 1 and 4 lineages in HOC and LOC lines, respectively.

### 2.4 Statistical analysis

Two mixed linear models, namely models 1 and 2, were exploited to estimate genetic parameters. Model 1 was as follows (Hong et al., 2013):

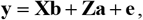

where **y** is the vector of phenotypic records; **b** is the vector of fixed effects; **a** is the vector of direct additive genetic effects; **e** is the vector of errors; **X** and **Z** is the design matrices. In this study, the data from the two lines were merged and then simultaneously analyzed. For OC and OCMBW, fixed effects were sex, generation × line, and days of age at measuring OC (linear and quadratic covariates). For BW traits, fixed effects were sex and generation × line. The mean and variance-covariance of the random effects were as follows:

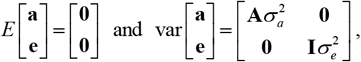

where 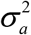 is the direct additive genetic variance; 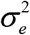 is the error variance; **A** is the additive relationship matrix; and **I** is the identity matrix. Variance components were estimated using ASREML software version 4.1 (Gilmour et al., 2015). Single-trait analysis was performed to estimate trait heritability, and two-trait analysis was performed to estimate genetic correlation between two traits. Genetic trends for OC, OCMBW, and BW8 were estimated using EBVs from the single-trait analysis.

Model 2 was as follows:

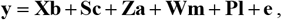

where **c** is the vector of the effects of maternal lineages (cytoplasmic inheritance) (e.g., Southwood et al., 1989; Grigoletto et al., 2017), **m** is the vector of maternal additive genetic effects; **l** is the vector of common litter environmental effects; and **S**, **W**, and **P** were the design matrices. The mean and variance-covariance of the random effects were as follows:

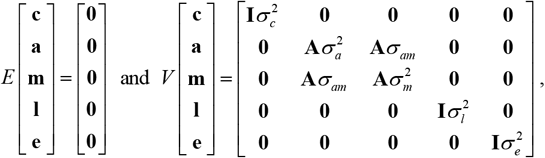

where 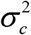 is the variance of cytoplasmic inheritance effects; 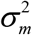 is the maternal additive genetic variance; *σ_am_* is the direct-maternal additive genetic covariance; and 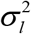 is the common litter environmental variance.

### 2.5 Resequencing genomic DNA from pools of mice in each strain

Genomic DNA extraction and resequencing were performed at the Macrogen Japan Co. Ltd. (Kyoto, Japan). Briefly, cryopreserved tail samples of 33 G16 mice used as parents of G17 in H line and 31 G16 mice used as parents of G17 in L line were available for DNA extraction. The DNA samples were prepared according to the Illumina TruSeq Nano DNA library preparation guide or TruSeq DNA PCR-free library preparation guide. The libraries were sequenced using Illumina NovaSeq 6000 sequencer. Each sequenced sample is prepared according to the Illumina TruSeq DNA sample preparation guide to obtain a final library of 300-400 bp average insert size. One microgram (TruSeq DNA PCR-free library) or 100 nanogram (TruSeq Nano DNA library) of genomic DNA is fragmented by covaris systems, which generates dsDNA fragments with 3’ or 5’ overhangs. Macrogen performed quality control analysis on the sample library and quantification of the DNA library templates. Read length was 150 base pairs (bp). The Ilumina Novaseq 6000 generated raw images and base calling through an integrated primary analysis software called RTA3 (Real Time Anarysis 3). The BCL (base calls) binary was converted into FASTQ using illumine package bcl2fastq2-v.20.0. The demultiplexing option (--barcode-mismatches) was set to perfect match (value: 0).

### 2.6 Variant calling and annotation

Variant calling analysis was performed for each of the 2 pooled DNA samples, one was from HOC line and the other was from LOC line, at the Macrogen Japan Co. Ltd. (Kyoto, Japan). Briefly, reads were mapped to mouse genome reference sequence (mm 10 from UCSC (Dec. 2011)) using Isaac Aligner v.01.15.02.08, designed to next-generation sequencing data with low-error rates, and then variants were called using Isaac Variant Caller v.2.0.13 (Raczy et al., 2013). Reference size was 2,652 Mbp and parameter settings were 15 for -base-quality cutoff, 1 for -keep-duplications, and AGCTCGGAAGAGC*, *GCTCTTCCGATCT for -default-adapters. Mappable yield and mean depth were 187,846 Mbp and 70.80 for pooled genomic DNA samples from H strain and 188,852 Mbp and 71.20 for those from L strain. Variants were annotated using SnpEff v.4.1 (Cingolani et al., 2012).

### 2.7 Selective signature detection

Fisher’s exact probability test for differences in allele frequencies was performed for the 4,147,082 autosomal variants commonly detected in the HOC and LOC lines (e.g., Guirao-Rico & González, 2021; Kofler et al., 2012; Lucotte et al., 2016). In this study, genomic regions with ≥5 consecutive variants with p-value lower than 0.05/4,147,085 (= 1.21 ×10^-8^) were defined as candidate regions, and genes tagged with the variants in the candidate regions were extracted as possible candidate OC-related genes.

## 3 RESULTS AND DISCUSSION

### 3.1 Genetic parameter estimation

Estimated heritability was 0.40 for OC 0.41 for OCMBW (Table 2). Hong et al. (2013) estimated heritability per line using phenotypic records of mice from G1 to G6 and from G1 to G9, and the estimated values ranged from 0.35 to 0.52 (0.48 in average) for OC and from 0.49 to 0.64 (0.55 in average) for OCMBW. The estimates in this study were slightly different from those of Hong et al. (2013), which might be due to the difference in the number of records and the fact that the data from the two lines were merged. Estimated heritabilities of BW traits were around 0.5. Hong et al. (2013) estimated heritability of BW8 to be ranging from 0.48 to 0.56 (0.51 in average), similar to our estimated value of 0.51.

**TABLE 2.** Phenotypic variance and direct heritability estimated using model 1

| **Item** | **OC** | **OCMBW** | **BW3** | **BW4** | **BW5** | **BW6** | **BW7** | **BW8** |
| --- | --- | --- | --- | --- | --- | --- | --- | --- |
| Phenotypic variance | 1,331.1 | 282,490 | 3.05 | 5.17 | 4.47 | 5.28 | 6.51 | 8.39 |
| SE | 38.56 | 8,290 | 0.08 | 0.14 | 0.13 | 0.15 | 0.19 | 0.27 |
| Heritability | 0.40 | 0.41 | 0.53 | 0.48 | 0.49 | 0.47 | 0.48 | 0.51 |
| SE | 0.03 | 0.03 | 0.02 | 0.03 | 0.03 | 0.03 | 0.03 | 0.03 |

SE: standard error. See Table 1 for other abbreviations.

Estimated genetic correlation between OC and OCMBW was 0.74 (Table 3). Hong et al. (2013) estimated the genetic correlation between OC and OCMBW, ranging from 0.77 to 0.94 (0.86 in average). A biological interpretation of the genetic correlation between OC and OCMBW lower than 1 could be that the main factors involved in OC are BW and body composition, whereas OCMBW might depend more on the proportion of the body occupied by internal organs with different metabolic rates (Stahl, 1965) and the metabolic rate at the cellular level, such as ATP production/resolution and mitochondrial proton leak capacity (Hulbert et al., 2002; Porter, 2001).

**TABLE 3.** Direct genetic correlation between traits estimated using model 1

| **Trait** | **OCMBW** | **BW3** | **BW4** | **BW5** | **BW6** | **BW7** | **BW8** |
| --- | --- | --- | --- | --- | --- | --- | --- |
| OC | 0.74 | 0.19 | 0.29 | 0.44 | 0.45 | 0.47 | 0.53 |
| SE | 0.03 | 0.06 | 0.06 | 0.06 | 0.06 | 0.06 | 0.06 |
| OCMBW | -- | -0.28 | -0.28 | -0.27 | -0.38 | -0.45 | -0.50 |
| SE | -- | 0.06 | 0.06 | 0.06 | 0.06 | 0.06 | 0.06 |
| BW8 | -- | 0.55 | 0.63 | 0.87 | 0.91 | 0.97 | -- |
| SE | -- | 0.04 | 0.04 | 0.02 | 0.02 | 0.01 | -- |

See Tables 1 and 2 for abbreviations.

Estimated genetic correlations of BW8 were 0.53 with OC and −0.50 with OCMBW (Table 3). Hong et al. (2013) estimated genetic correlation between OC and BW8 to be ranging from 0.07 to 0.50 (0.22 in average), and that between OCMBW and BW8 to be ranging from −0.51 to −0.03 (−0.27 in average). In this population, there were periods when the indicator traits for selection differed between lines, due to that the purpose of the selection experiment was to produce two mice lines with different OC without changing BW and that correlated responses in BW were observed (Figures 2, 3, Table 3). In such cases, restricted selection on OC and BW might be more effective (Kempthorne & Nordskog, 1959; Satoh, 2019; Yamada et al., 1975).

The absolute values of the estimated genetic correlations of BW traits with OC and OCMBW were larger when the age at measuring BW was closer to that at measuring OC (Table 3). Using an animal model with maternal effects, Leamy et al. (2005) estimated genetic correlations of heat loss measured at 13 weeks of age in average 3 to be 0.18 with BW at 3 weeks (weaning), 0.24 with BW with 6 weeks, and 0.36 with BW at 12 weeks.

### 3.2 Impact of maternal and cytoplasmic inheritance effects on genetic parameter estimation

For all traits, estimated heritabilities by using model 2 were lower and their standard errors were larger than those obtained using model 1 (Tables 3, 4). Previous studies have also reported that adding of maternal effects increased the standard errors (e.g., Meyer et al., 1992; Ogawa et al., 2021; Solanes et al., 2004). Estimated direct heritabilities were 0.32 for both OC and OCMBW. Leamy et al. (2005) estimated the direct heritability of heat loss in mice to be 0.20 using an animal model considering common litter environmental effects. Estimated direct heritabilities of BW were lower at younger ages, around 0.1 or lower for BW3, BW4, BW5, and BW6, and around 0.2 for BW7 and BW8. Estimated residual variances in BWs were almost unchanged between the models, except for BW3. In simulation studies, direct heritability of trait affected by maternal effects has been reported to be overestimated when using animal models ignoring the maternal effect (Clément et al., 2001; Satoh et al., 2002; Southwood et al., 1989).

**TABLE 4.** Genetic parameters estimated using model 2

| Item | OC | OCMBW | BW3 | BW4 | BW5 | BW6 | BW7 | BW8 |
| --- | --- | --- | --- | --- | --- | --- | --- | --- |
| Phenotypic variance | 1,293.7 | 273,060 | 3.36 | 5.19 | 4.48 | 5.31 | 6.62 | 8.39 |
| SE | 39.0 | 8,795 | 0.16 | 0.20 | 0.17 | 0.19 | 0.25 | 0.33 |
| Direct heritability | 0.32 | 0.32 | 0.08 | 0.06 | 0.09 | 0.12 | 0.20 | 0.22 |
| SE | 0.06 | 0.07 | 0.06 | 0.06 | 0.06 | 0.06 | 0.06 | 0.07 |
| Maternal heritability | 0.02 | 0.03 | 0.13 | 0.11 | 0.07 | 0.06 | 0.07 | 0.06 |
| SE | 0.04 | 0.04 | 0.07 | 0.06 | 0.05 | 0.05 | 0.05 | 0.05 |
| Direct-maternal genetic correlation | −0.79 | −0.96 | 0.26 | 0.14 | 0.63 | 0.87 | 0.39 | 0.38 |
| SE | 0.46 | 0.49 | 0.51 | 0.54 | 0.81 | 0.86 | 0.46 | 0.52 |
| Proportion of common litter environmental variance | 0.12 | 0.13 | 0.52 | 0.33 | 0.29 | 0.21 | 0.16 | 0.15 |
| SE | 0.03 | 0.03 | 0.06 | 0.05 | 0.04 | 0.04 | 0.03 | 0.04 |
| Proportion of variance of cytoplasmic inheritance effects | 0.01 | 0.02 | 0.00 | 0.01 | 0.00 | 0.00 | 0.01 | 0.00 |
| SE | 0.01 | 0.02 | 0.00 | 0.01 | 0.01 | 0.01 | 0.02 | 0.01 |

2 See Tables 1 and 2 for abbreviations.

Estimated maternal heritabilities were about 0.1 or lower for all traits and smaller than twice the standard errors (Table 4). Direct-maternal genetic correlations were estimated to be negative for OC and OCMBW and positive for BW traits, but their standard errors were large. These results suggest that there is little need to consider maternal genetic effects. Accurate genetic parameter estimation requires a sufficient number of records, as well as an appropriate model (Thompson, 1976) and data structure (Gerstmayr, 1992). It has been argued that negative estimates of direct-maternal genetic correlations might be artifacts (e.g., Beniwal et al., 1992; Bijma, 2006; Lee, 2001; Willham, 1980). Recently, the possible involvement of genomic imprinting has also been pointed out (Hager et al., 2008; Meyer & Tier, 2012; Varona et al., 2015). These facts require careful interpretation of the results.

Estimated values of the proportion of common litter environmental variance to the total phenotypic variance were around 0.1 for OC and OCMBW, whereas the value was higher at younger ages for BW traits, 0.52 for BW3 compared to 0.15 for BW8 (Table 4). Formoso-Rafferty et al. (2016) estimated maternal heritability and the proportion of common litter environmental variance of weaning weight (BW3) to be 0.16 and 0.41, respectively. Estimates of the maternal additive variance in BW traits did not largely change between different weeks, those of permanent environmental variance tended to decrease with increasing age, and those of the direct additive genetic and error variances increased with increasing age. This was likely to be the main cause of the decrease in maternal heritability and the proportion of common litter environmental variance with increasing age. Two-trait animal model analysis including all the random effects did not converge, and thus, two-trait animal model analysis including only common litter environmental effects was performed. As a result, estimated common litter environmental correlations of BW8 were 0.69, 0.77, 0.92, 0.92, 0.95, and 0.68 with BW3, BW4, BW5, BW6, BW7, and OC. These results suggest that the common litter environmental effects on weaning weight (BW3) could be highly persistent and that the common litter environmental effects on BW traits might indirectly affect OC and OCMBW measured after 8 weeks. Previous studies suggested the possibility that although maternal effects on BW in mice tends to weaken with increasing weeks of age, the effects might persist after maturation (e.g., Cowley et al., 1989; Leamy et al., 2005). Beniwa1 et al. (1992) estimated the proportion of common litter environmental variance of BW at 10 weeks in mice and discussed the possibility that the results were caused by the fact that all same-sex littermates were kept in the same cage after weaning. In this population, full-sibs of the same sex were kept in the same cage after weaning from G0 to G10, and therefore, there might be possible confounding of the effects of rearing environments before and after weaning on BW traits. In addition, the number of pups was adjusted to 6 in principle, but some exceptions were observed. Data were also analyzed after excluding such exceptions, and the results were similar with those obtained when the data were not excluded (results not shown). Therefore, it was considered unlikely that differences in the number of pups were not the main factor for the common litter environmental effects in this study.

Estimated values of the proportion of the variance of the cytoplasmic inheritance effects, regarding as mitochondrial DNA, were very low for all traits (Table 4). Also, adding the effects of maternal lineages were considered as the fixed effect in model 1 did not affect the genetic parameter estimation (results not shown). These results suggest that there is no need to consider the cytoplasmic genetic effects in this study. Possible reasons are 1) that the effect of mitochondrial inheritance is truly weak, 2) that the diversity of mitochondrial DNA in the F0 population might be already low in this population (Petters et al.,1988), 3) cytoplasmic inheritance effects might also consider factors other than mitochondria, and 4) discrepancies between points related to inter- and intra-individual transmission of mitochondrial DNA and assumptions about cytoplasmic inheritance effects such as mitochondrial DNA heteroplasmy (e.g., Stewart & Chinnery, 2015). With respect to 1), Southwood et al. (1989) reported that when analyzing simulated data with maternal genetic effects but without cytoplasmic inheritance effects with an animal model considering both effects, a small value was estimated as the proportion of variance of cytoplasmic inheritance effects. However, it is difficult to determine whether the estimated small values are true or not unless the true genetic model is known for real data analysis (Southwood et al., 1989). For 4), not only oxidative stress-induced damage of mitochondrial DNA but also initialization mechanism of mitochondrial DNA (Ling et al., 2016) might be involved, but such information is not considered in the covariance structure for cytoplasmic inheritance effects in this study (Southwood et al., 1989). It might be important to investigate the approach as in Quintanilla et al. (1999), where the correlation of the effects among individuals belonging to the same maternal lineage is assumed to be <1. Furthermore, the relationship between mitochondrial DNA and phenotype is still unclear (Ghiselli & Milani, 2020), and further study is needed.

### 3.3 Possible candidate causal genes for oxygen consumption

According to the moderate direct heritability and negligible proportion of the variance of cytoplasmic inheritance effects (Table 4), we speculated that the genetic causes of the difference in OC between lines are likely to exist on the nuclear genome. In order to test this hypothesis as much as possible, we conducted a selective signature analysis using the results of pool-seq. Accumulating insight into genes relating to OC not only could contribute to develop easier approach for estimating MER and RFI (e.g., Cantalapiedra-Hijar et al., 2018; Cooper-Prado et al., 2014; Gilbert et al., 2017), but also might give interesting information in terms of elucidating nuclear-mitochondrial crosstalk (e.g., Cannino et al., 2007; Liu et al., 2017; Poyton & McEwen, 1996).

There are several indicators proposed for detecting selective signatures (Cadzow et al., 2014; Ma et al., 2015). Initially, we planned to use the indicator ZHp proposed by Rubin et al. (2010), but the histogram of ZHp was far from a bell-shaped distribution. Therefore, we decided to implement Fisher’s exact probability test in this study. Fisher’s exact probability test has been widely used in previous studies on various species and populations (e.g., Guirao-Rico & González, 2021; Horns et al., 2019; Kofler et al., 2012; Lucotte et al., 2016; Riva et al, 2020; Shapiro & Alm, 2008). The Manhattan plots per autosome are shown in Figures S1 through S19. We adopted the Bonferroni-corrected value (0.05/4,147,085 = 1.2×10^-8^) as a conservative threshold, but aiming to further reduce false positives, genomic regions with ≥5 consecutive variants with p-values lower than the threshold were detected as selective signatures. We finally extracted the 2,792 genes as possible candidate causal genes relating to OC (Table S1).

Information on the extracted genes is important for understanding the biological mechanisms of OC and correlated responses observed in the selection experiments (Darhan et al., 2017, 2019; Hong et al., 2015). The extracted 2,792 genes included, for example, *Ndufa12* gene which is thought to be related to complex I controlling the electron transfer system (e.g., Rak & Rustin, 2014); *Sdhc* gene which is thought to be related to complex II (e.g., Ishii et al., 2005); *Uqcc3* gene which is thought to be related to complex III (e.g., Yang et al., 2020), and *Atp10b* gene related to ATP synthase (e.g., Smolders & Broeckhoven, 2020). These results might explain the differences in hepatic mitochondrial respiratory rate and proton leak level between HOC and LOC lines found in previous studies (Darhan et al., 2019; Hong et al., 2015).

Genes suspected to be associated to feed efficiency were extracted, including *Fto* gene which is known to be related to obesity (e.g., Church et al., 2009) *Stat6* gene which is thought to be related to lipid production (e.g., Ricardo-Gonzalez et al., 2010), and *Phospho1* gene encoding enzyme possibly regulating thermogenesis, insulin resistance, and obesity (e.g., Jiang et al., 2020; Suchacki et al., 2020). Genes previously reported as candidate ones for feed efficiency and related traits were also included, such as *Etv4* (e.g., Brunes et al., 2020; Weber et al., 2016), *Fgd6* (e.g., Ghoreishifar et al., 2020), *Fgf10* (e.g., Miao et al., 2017), *Ghsr* (e.g., Jin et al., 2014), and *IGF2BP1* genes (e.g., Chen et al., 2019; Liu et al., 2021). These findings might support the difference in chemical components, feed efficiency, and MER between the lines reported by Darhan et al. (2017).

Furthermore, *Actn3* gene, which is known to be expressed in fast twitch muscle (e.g., Niemi & Majamaa, 2005), was extracted, suggesting that the proportion of fast twitch versus slow twitch muscle in skeletal muscle might differ between the lines. On the other hand, genes for interleukin and Tumor Necrosis Factor (TNF) systems, which is known to be related to inflammatory cytokines, were extracted, suggesting interesting possibility of the relationship between OC and crosstalk between mitochondria and immune function (e.g., Angajala et al., 2018; Oliva-Ramírez et al., 2014; Qualls et al., 2021; Riley & Tait, 2020). In addition, *Apod* gene encoding Apolipoprotein D possibly relating to oxidative stress and aging (e.g., Dassati et al., 2014), *Prr16* gene encoding Largen protein (e.g., Yamamoto et al., 2014), and *Cry1* gene encoding a key component of the circadian core oscillator (e.g., Dimova et al., 2019) were extracted.

On the other hand, genetic selection has been carried out successively in closed populations based on EBV calculated assuming the infinitesimal model (Barton et al., 2017; Bulmer, 1980; Fisher, 1919), the population size is not large, and the base population was derived from four inbred lines (Figure 1; Hong et al., 2013, 2015). Therefore, false positives were not likely to be completely removed. Also, the results obtained might include effects related to cold stress tolerance (e.g., Hankenson et al., 2018; Hylander et al., 2019; Vialard & Olivier, 2020), because the population studied was reared at 22°C to 24°C. To detect candidate genes with higher resolution, it would be necessary to perform the individual-level sequencing.

## 4 CONCLUSIONS

We estimated genetic parameters using an animal model including maternal genetic, common litter environmental, and cytoplasmic inheritance effects for OC, OCMBW, and BW at 3 through 8 weeks of age in mice. Direct heritabilities of both OC and OCMBW were estimated to be both moderate, and the proportions of the variance of cytoplasmic inheritance effects were very low. We searched selective signatures on autosomes by performing pool-seq analysis. A list of possible candidate causal genes for OC was provided, including those relating to electron transport chain and ATP-binging proteins. We believe that the results could contribute to elucidate the genetic mechanism of OC, an indicator of MER.

## ACKNOWLEDGEMENTS

This research was partly supported by the Grant-in-Aid (18K14565) of the Japan Society for the Promotion of Science (JSPS), Japan. We thank Akiko Iwai, a technical assistant the Laboratory of Animal Breeding and Genetics, Graduate School of Agricultural Science, Tohoku University, for her help in preparing cryopreserved tail samples of OC mice for genome DNA extraction. We are grateful to all technical assistants and students of the Laboratory of Animal Breeding and Genetics, Graduate School of Agricultural Science, Tohoku University for their help in rearing and phenotypic data collection. Thanks go to all technical staffs of the Animal experiment building of the Graduate School of Agricultural Science, Tohoku University, for maintaining the rearing room environments.

## DATA AVAILABILITY STATEMENT

A request to the row data might be sent to the corresponding author.

## AUTHORS’ CONTRIBUTION STATEMENT

Shinichiro Ogawa: Conceptualization, Format analysis, Funding acquisition, Investigation, Methodology, Software, Visualization, Writing - original draft. Hongyu Darhan: Data curation, Resources, Writing - review & editing. Keiichi Suzuki: Conceptualization, Data curation, Methodology, Resources, Writing - review & editing.

