## Supplemental Figures 1 to 19 for "Genetic and genomic analysis of oxygen consumption in mice"

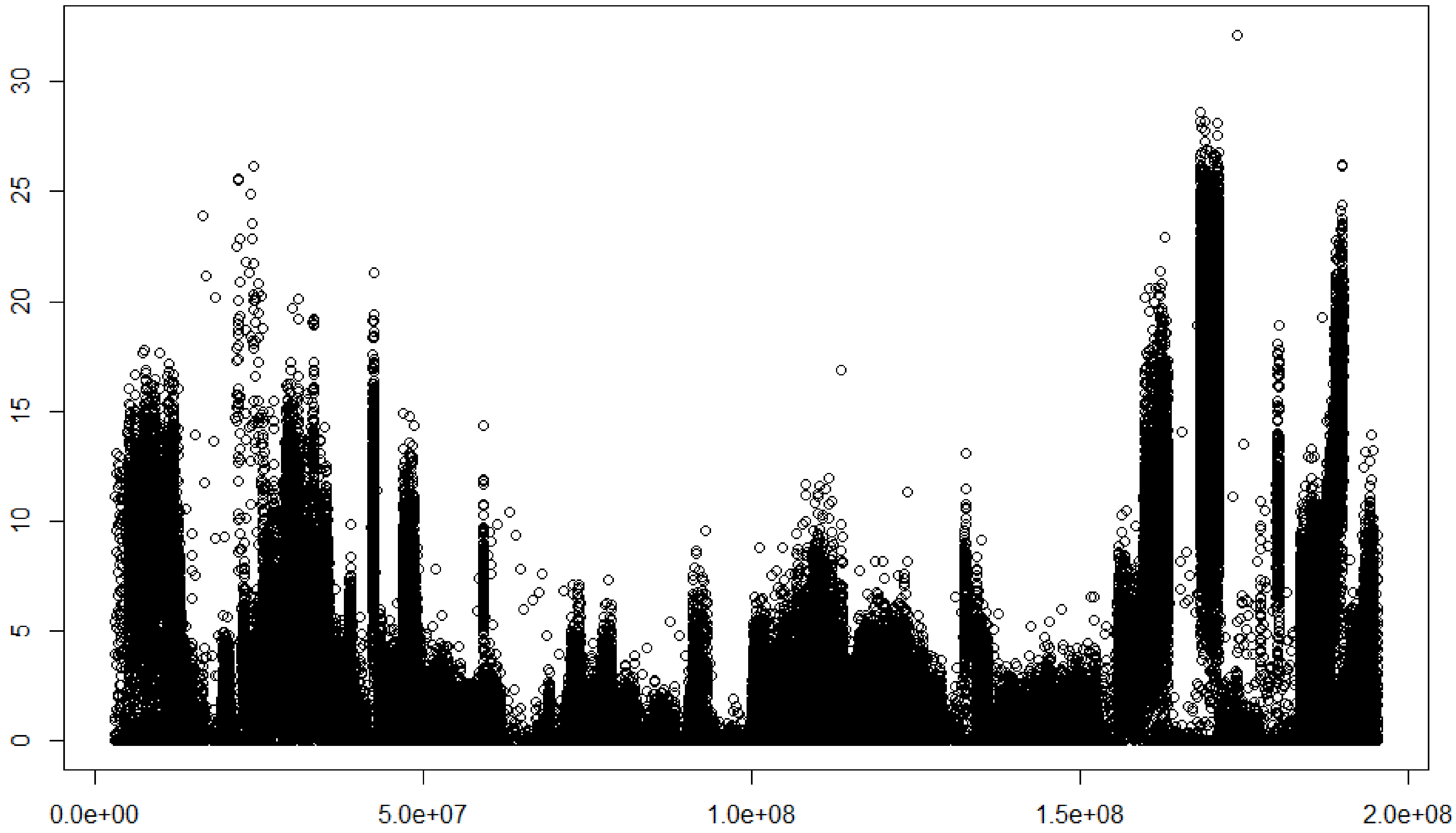


**Figure S1**. Manhattan plot of the results from fisher's exact test for difference in allele frequency of 309,245 variants on Mouse chromosome 1. Horizontal axis, location (base pair); Vertical axis, -ln(p) value. Threshold value for significance = -ln(0.05/4,147,085) = 18.23.

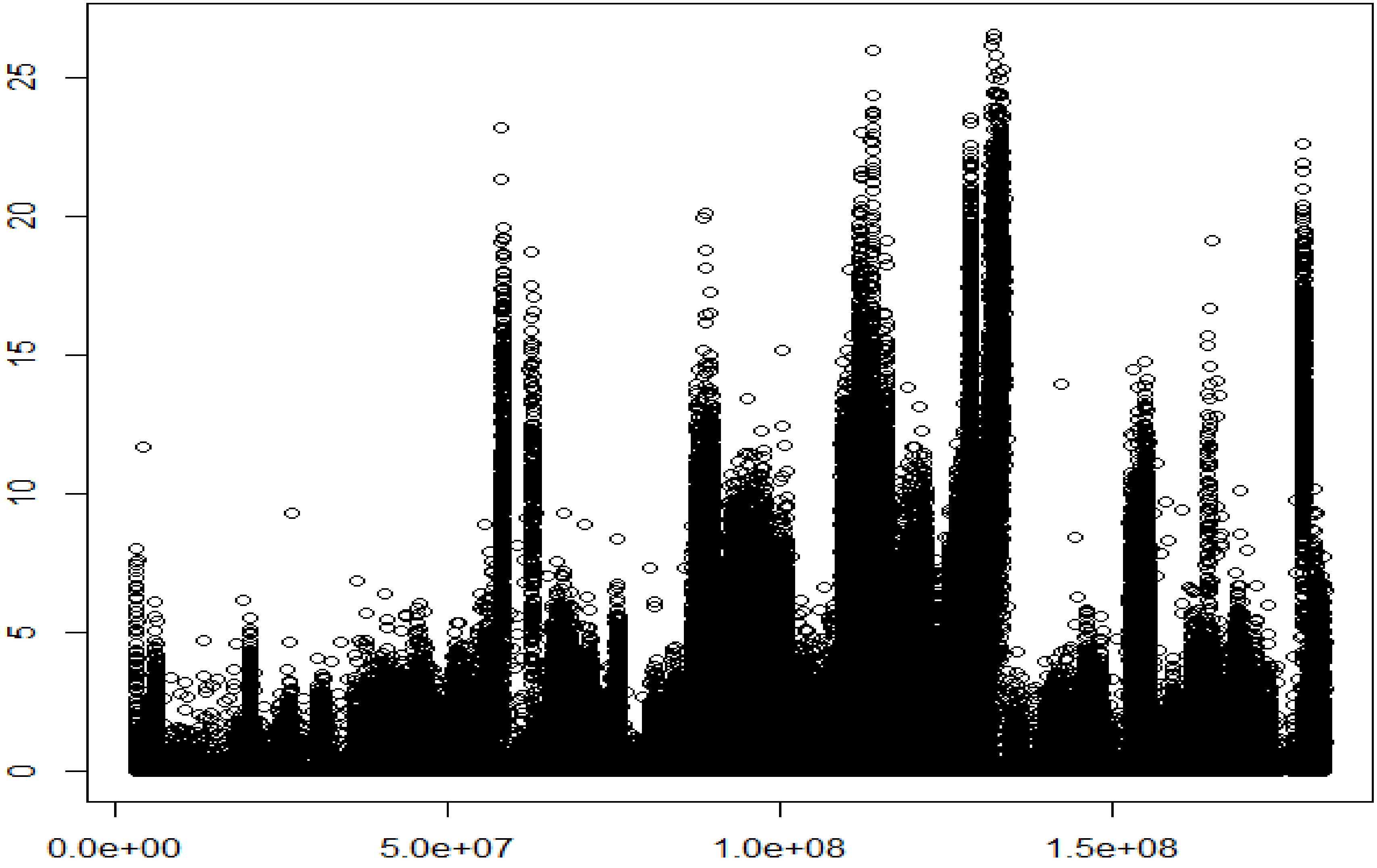


**Figure S2**. Manhattan plot of the results from fisher's exact test for difference in allele frequency of 258,563 variants on Mouse chromosome 2. Horizontal axis, location (base pair); Vertical axis, -ln(p) value. Threshold value for significance = -ln(0.05/4,147,085) = 18.23.

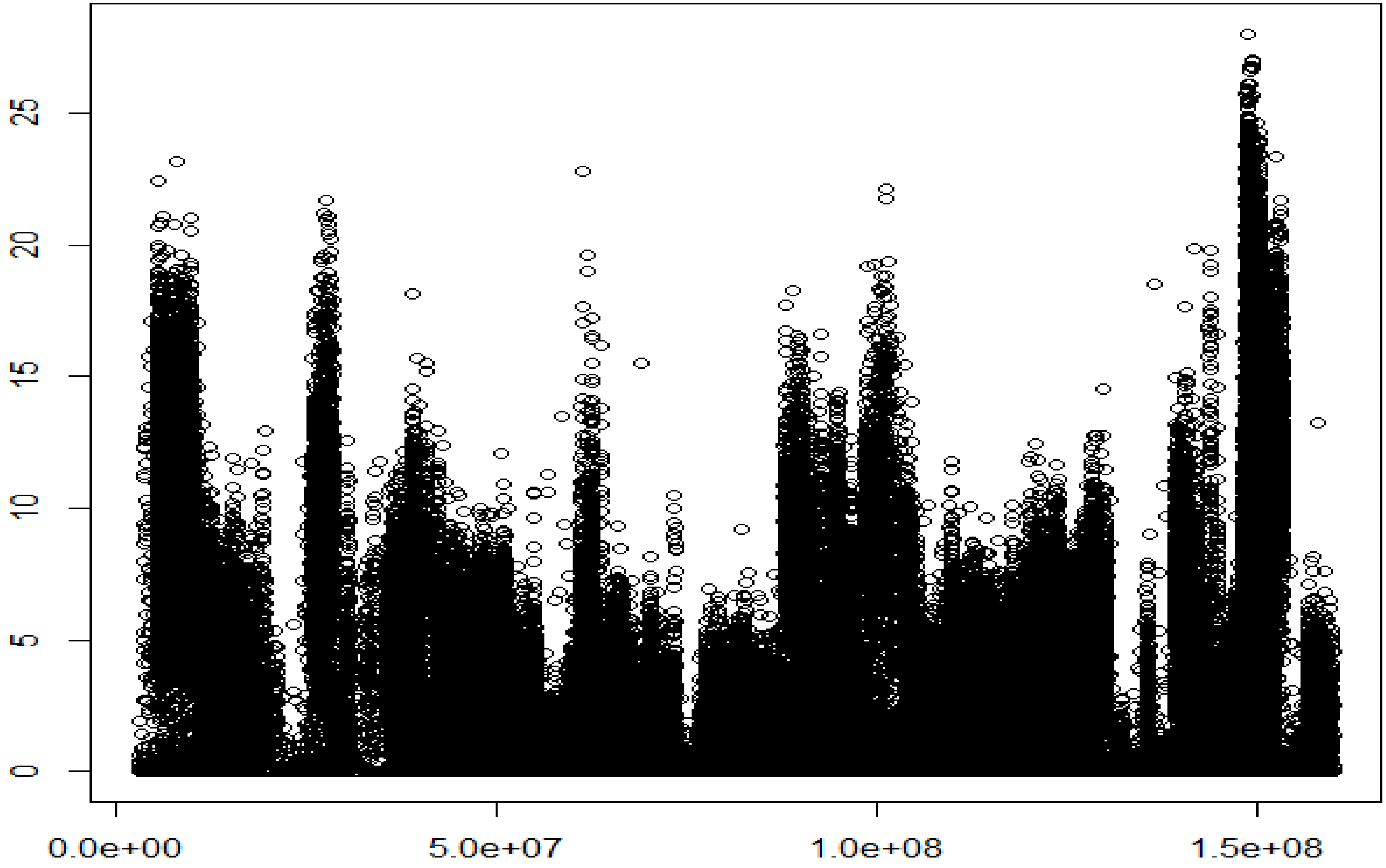


**Figure S3**. Manhattan plot of the results from fisher's exact test for difference in allele frequency of 294,418 variants on Mouse chromosome 3. Horizontal axis, location (base pair); Vertical axis, -ln(p) value. Threshold value for significance = -ln(0.05/4,147,085) = 18.23.

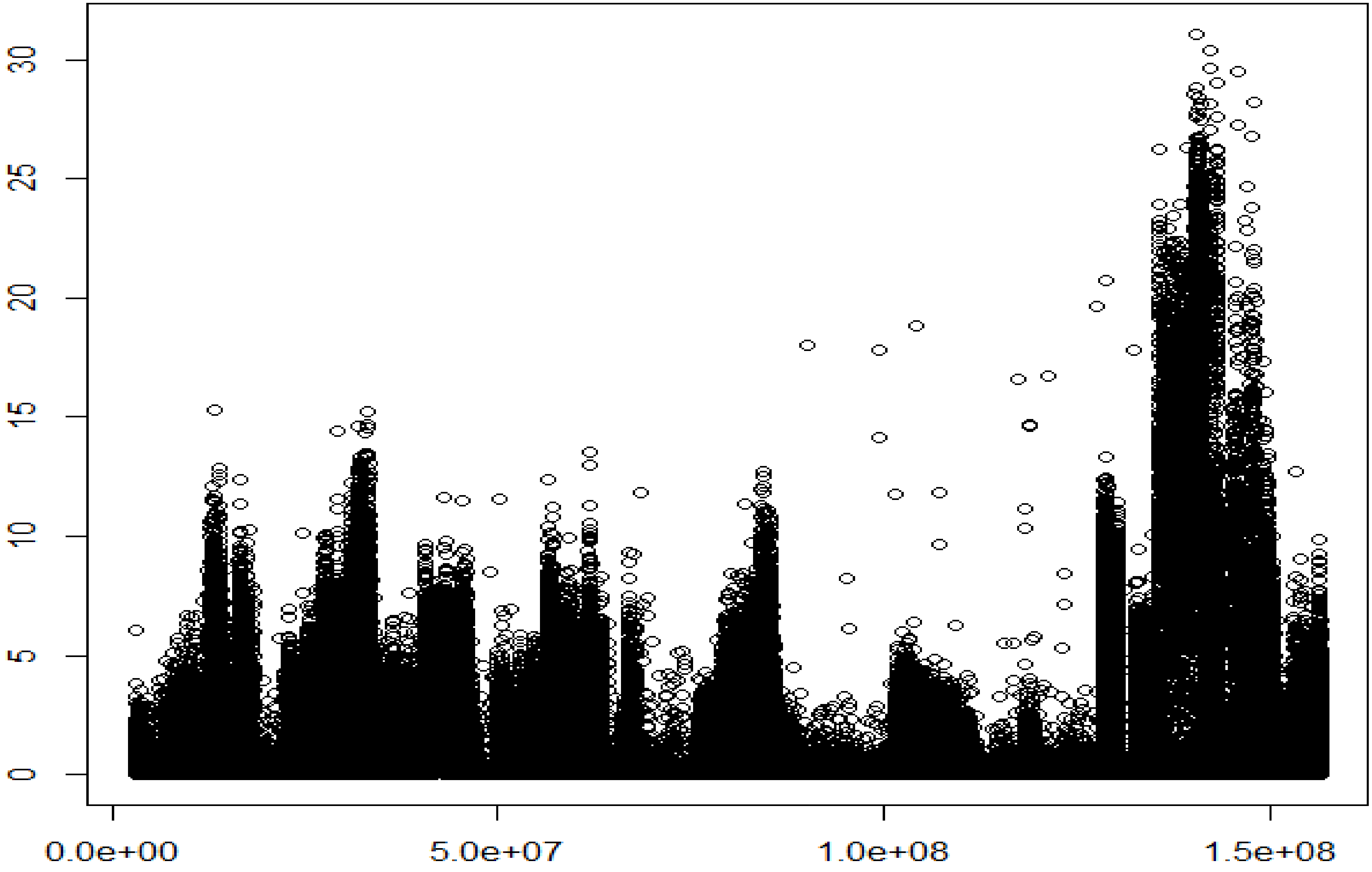


**Figure S4**. Manhattan plot of the results from fisher’s exact test for difference in allele frequency of 266,382 variants on Mouse chromosome 4. Horizontal axis, location (base pair); Vertical axis, -ln(p) value. Threshold value for significance = -ln(0.05/4,147,085) = 18.23.

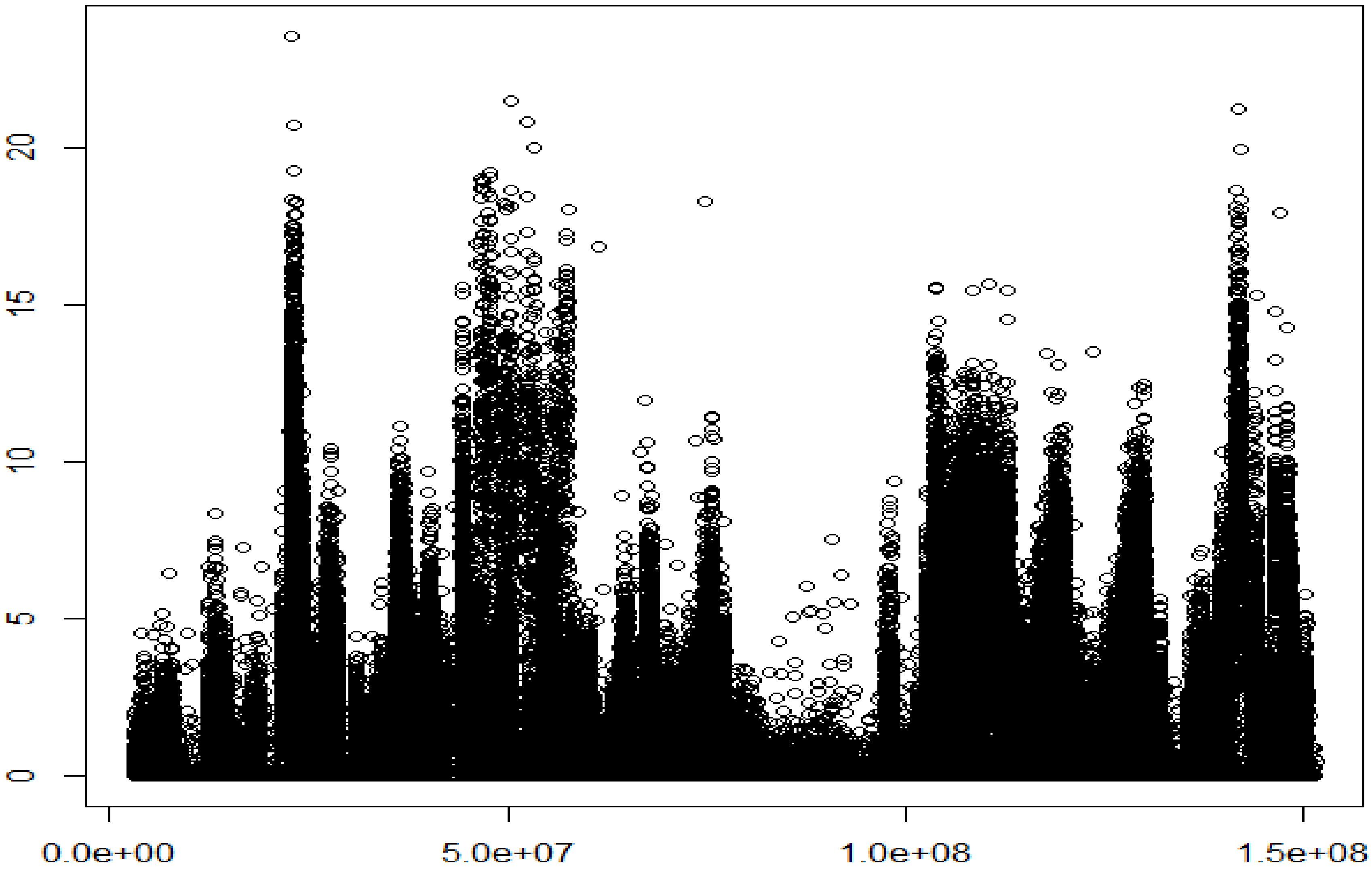


**Figure S5**. Manhattan plot of the results from fisher's exact test for difference in allele frequency of 255,151 variants on Mouse chromosome 5. Horizontal axis, location (base pair); Vertical axis, -ln(p) value. Threshold value for significance = -ln(0.05/4,147,085) = 18.23.

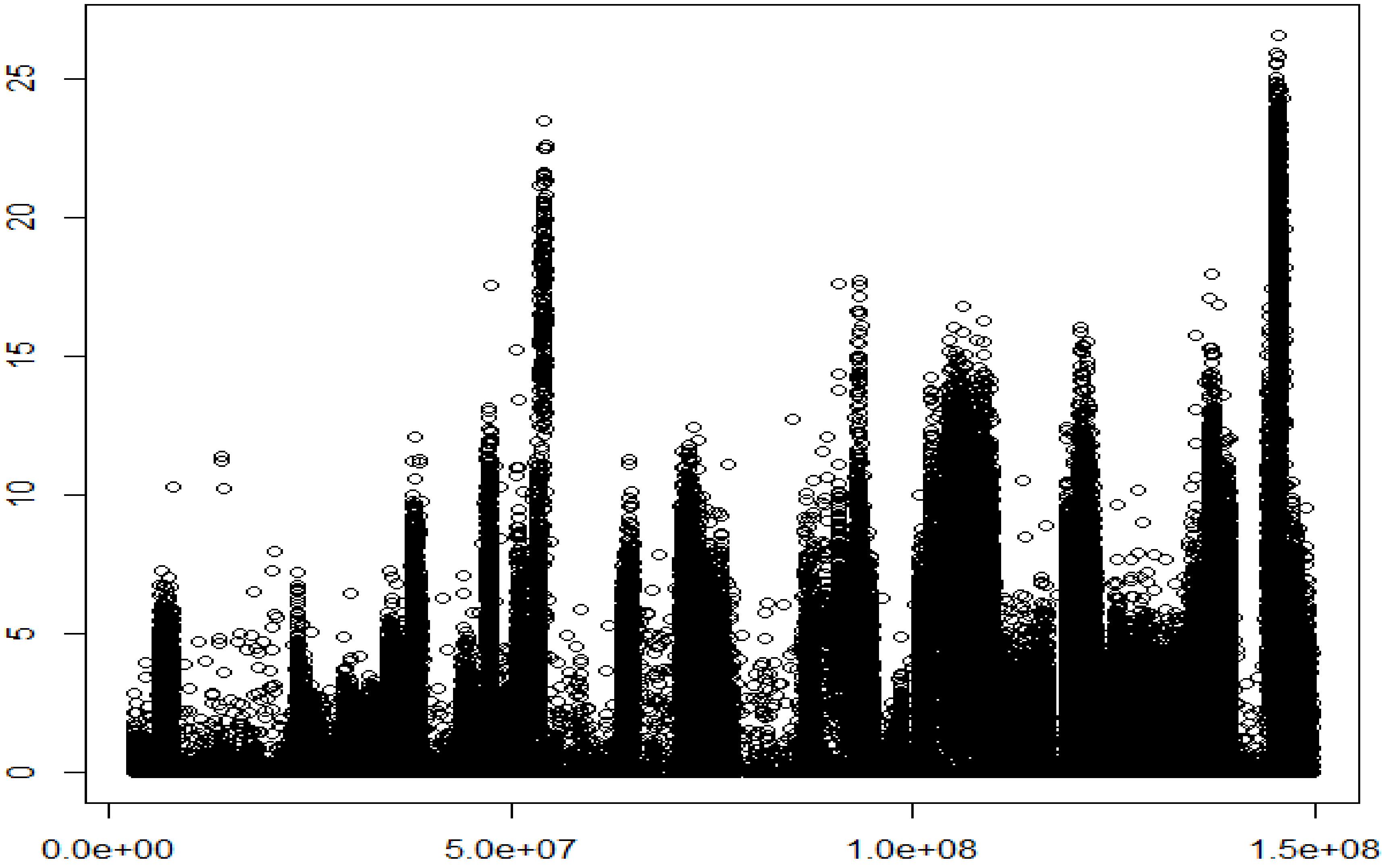


**Figure S6**. Manhattan plot of the results from fisher's exact test for difference in allele frequency of 235,535 variants on Mouse chromosome 6. Horizontal axis, location (base pair); Vertical axis, -ln(p) value. Threshold value for significance = -ln(0.05/4,147,085) = 18.23.

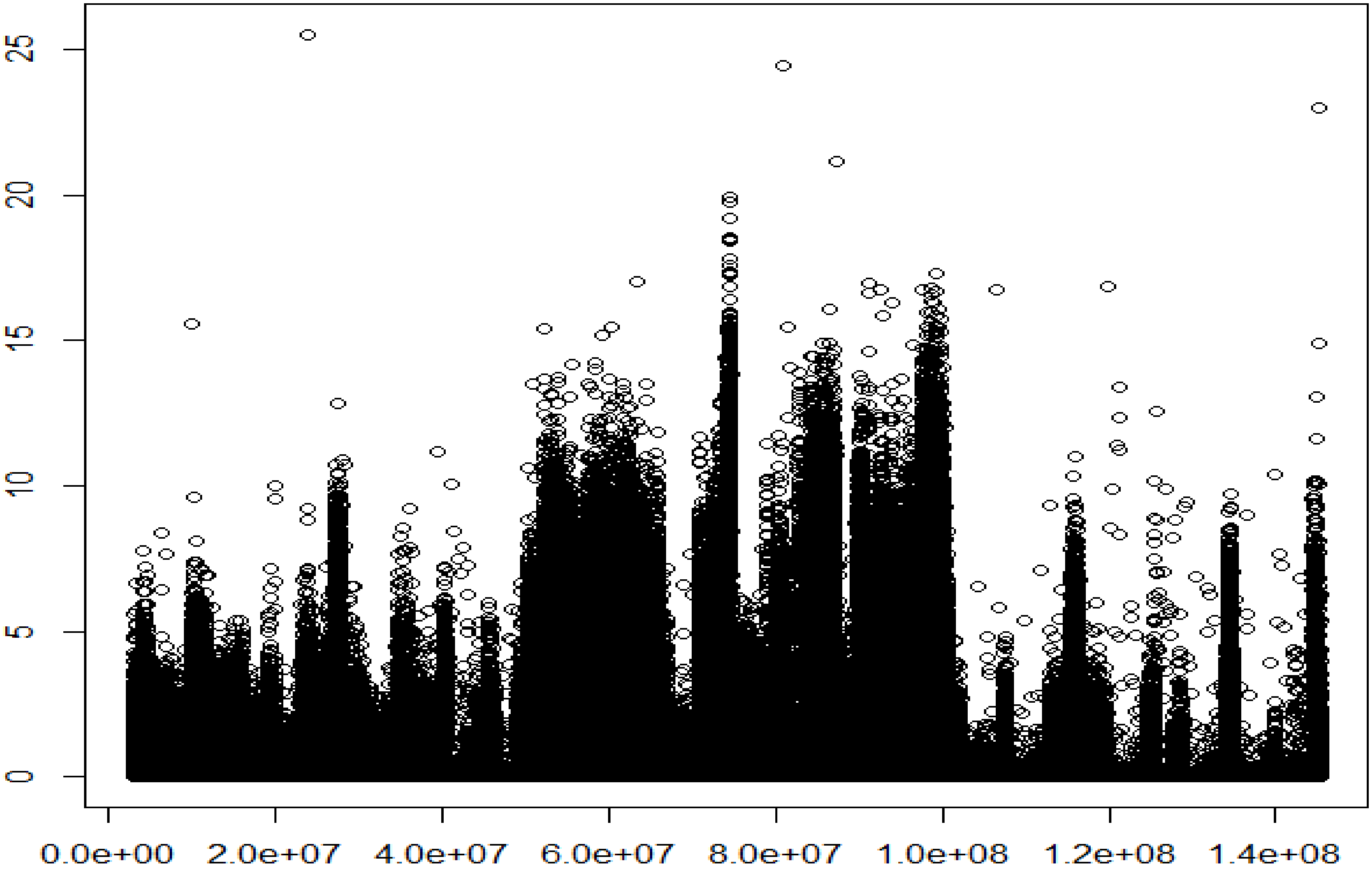


**Figure S7**. Manhattan plot of the results from fisher's exact test for difference in allele frequency of 300,694 variants on Mouse chromosome 7. Horizontal axis, location (base pair); Vertical axis, -ln(p) value. Threshold value for significance = -ln(0.05/4,147,085) = 18.23.

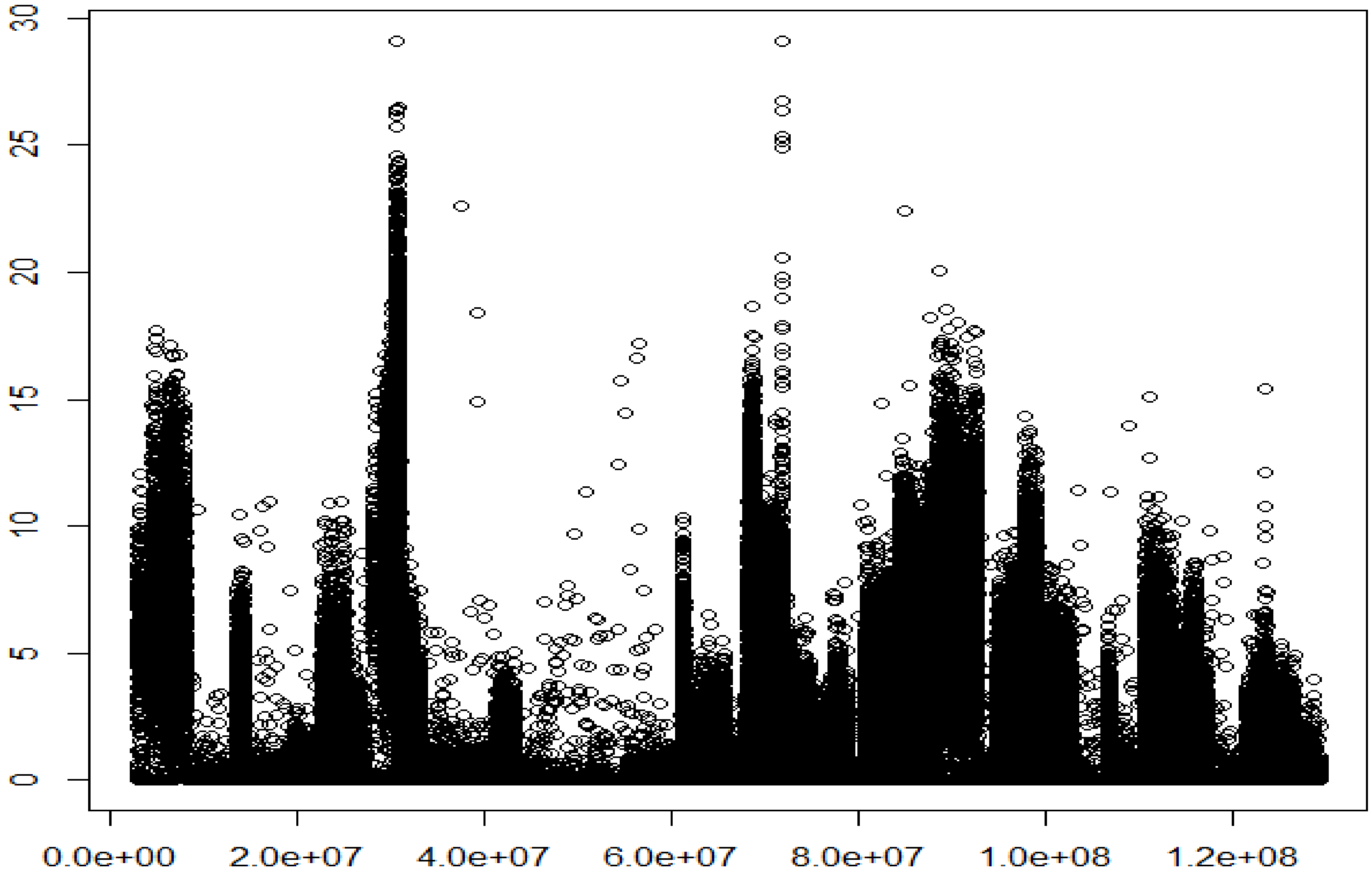


**Figure S8**. Manhattan plot of the results from fisher’s exact test for difference in allele frequency of 352,136 variants on Mouse chromosome 8. Horizontal axis, location (base pair); Vertical axis, -ln(p) value. Threshold value for significance = -ln(0.05/4,147,085) = 18.23.

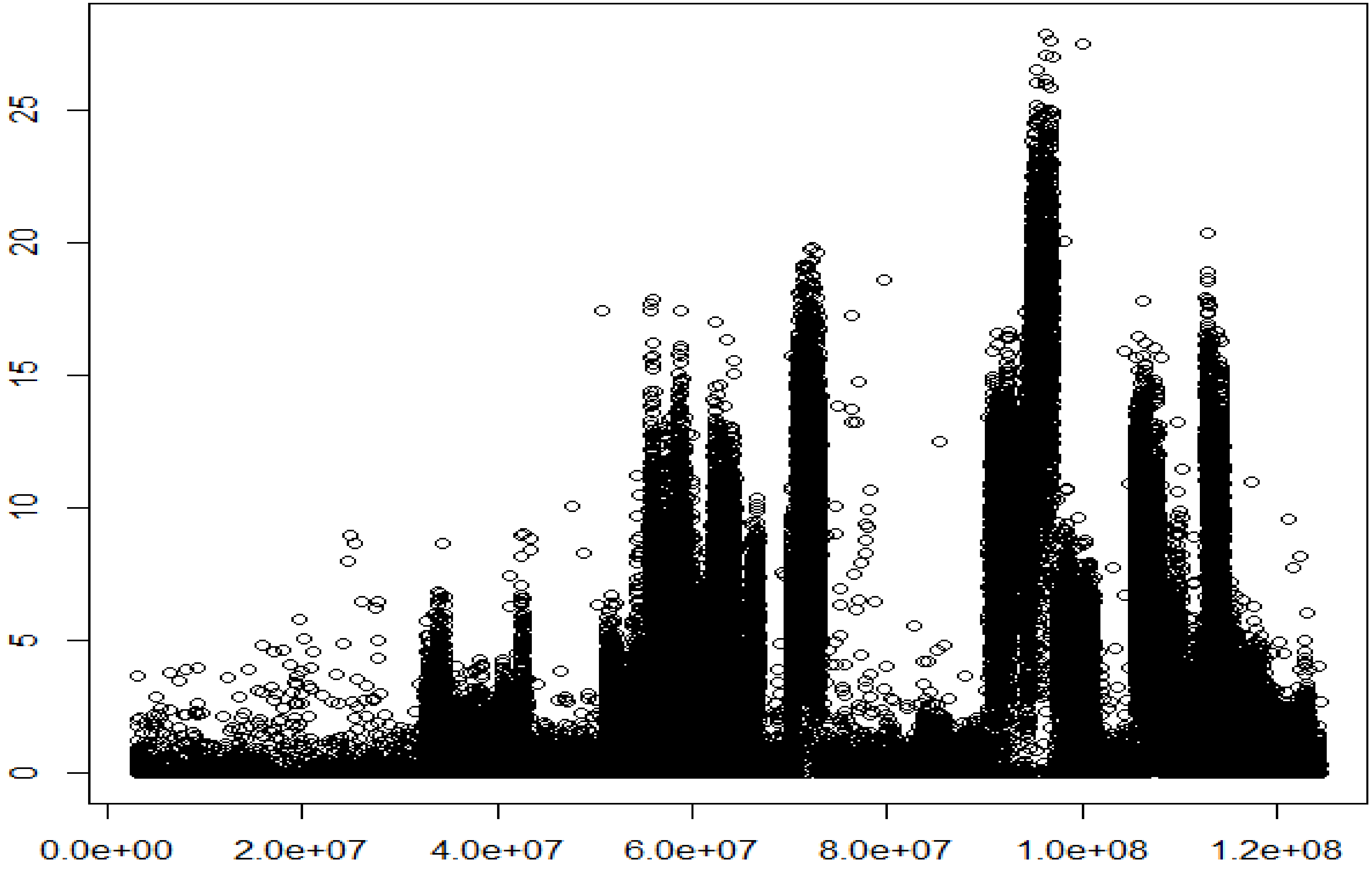


**Figure S9**. Manhattan plot of the results from fisher's exact test for difference in allele frequency of 188,961 variants on Mouse chromosome 9. Horizontal axis, location (base pair); Vertical axis, -ln(p) value. Threshold value for significance = -ln(0.05/4,147,085) = 18.23.

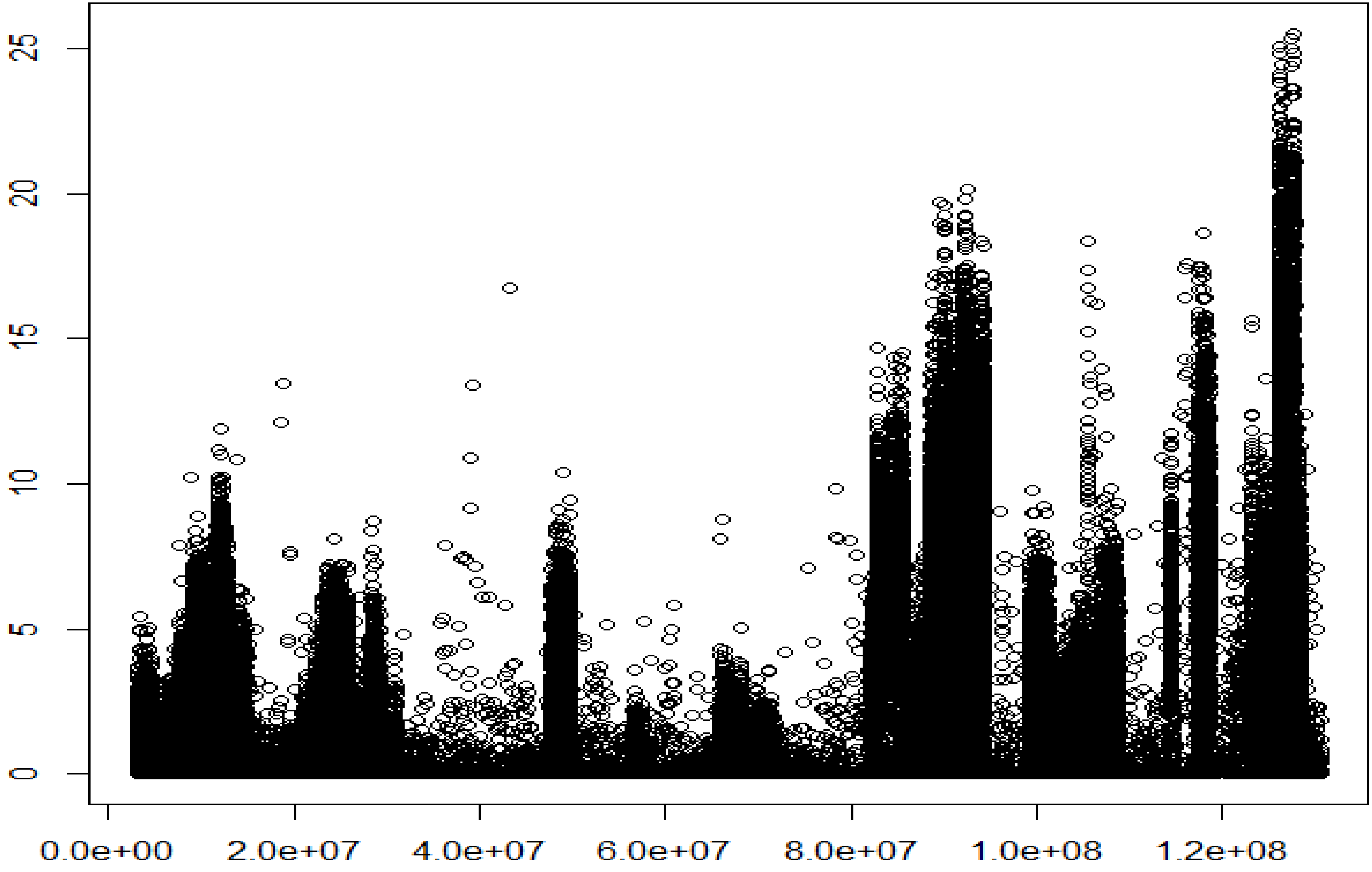


**Figure S10**. Manhattan plot of the results from fisher's exact test for difference in allele frequency of 133,197 variants on Mouse chromosome 10. Horizontal axis, location (base pair); Vertical axis, -ln(p) value. Threshold value for significance = -ln(0.05/4,147,085) = 18.23.

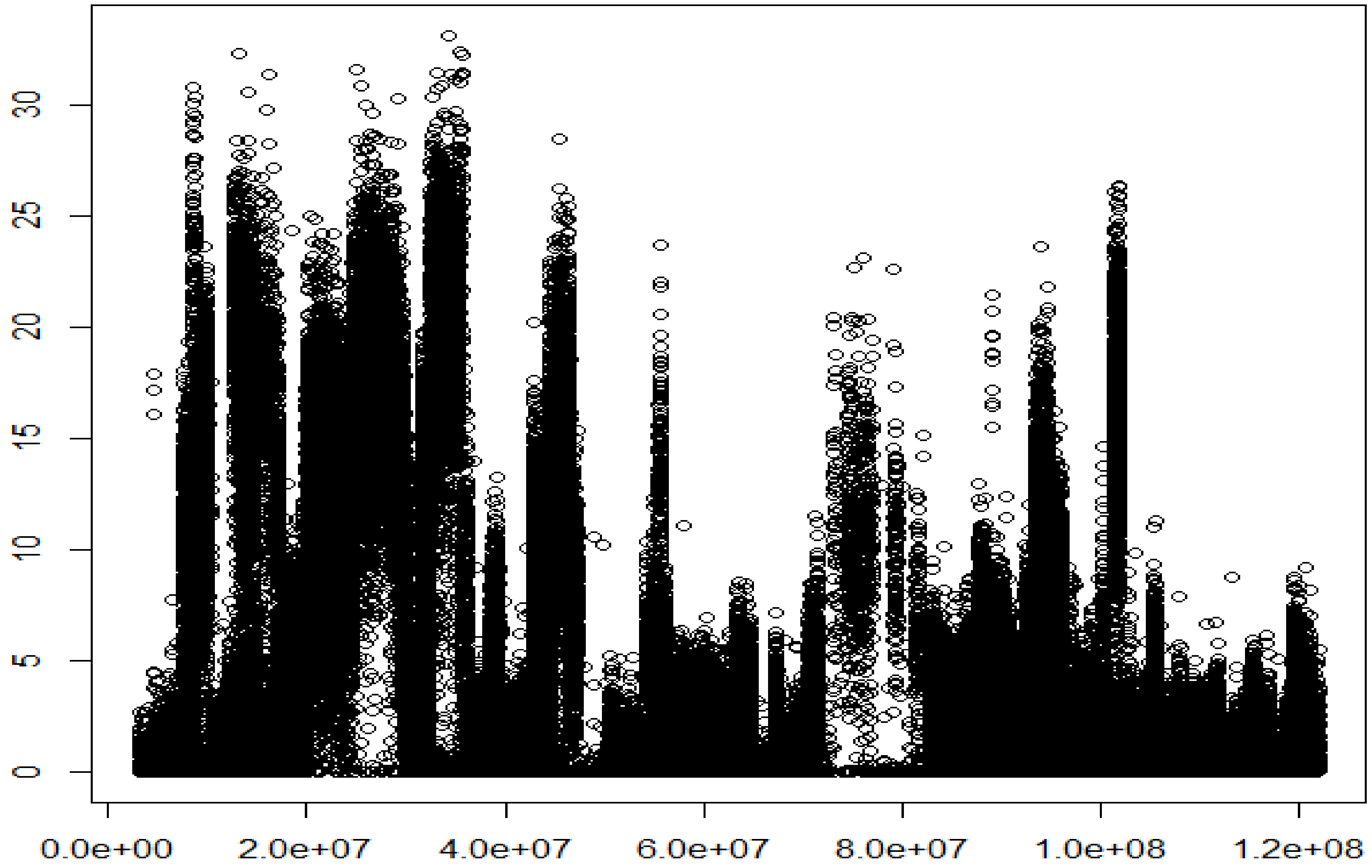


**Figure S11**. Manhattan plot of the results from fisher's exact test for difference in allele frequency of 226,696 variants on Mouse chromosome 11. Horizontal axis, location (base pair); Vertical axis, -ln(p) value. Threshold value for significance = -ln(0.05/4,147,085) = 18.23.

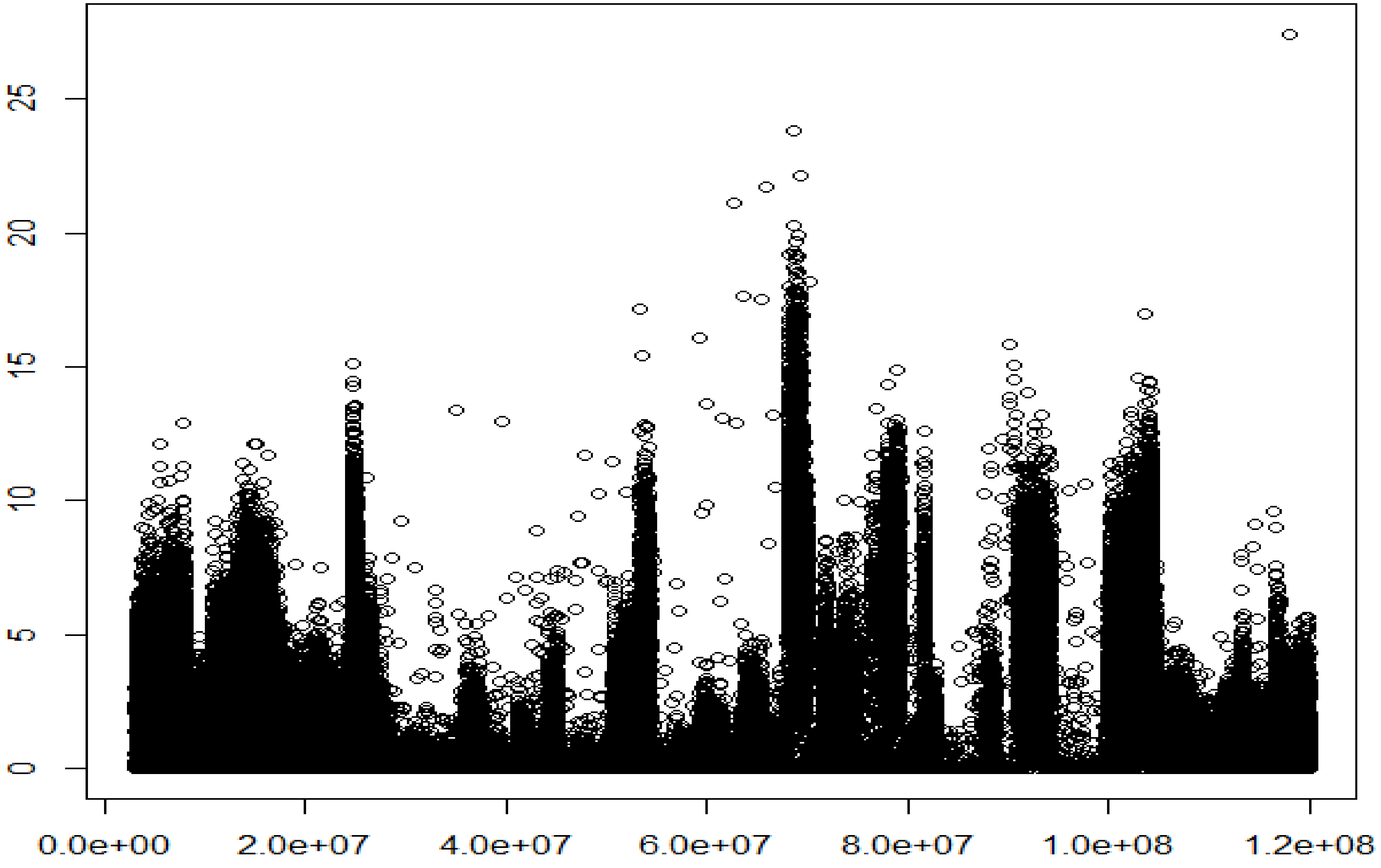


**Figure S12**. Manhattan plot of the results from fisher's exact test for difference in allele frequency of 277,395 variants on Mouse chromosome 12. Horizontal axis, location (base pair); Vertical axis, -ln(p) value. Threshold value for significance = -ln(0.05/4,147,085) = 18.23.

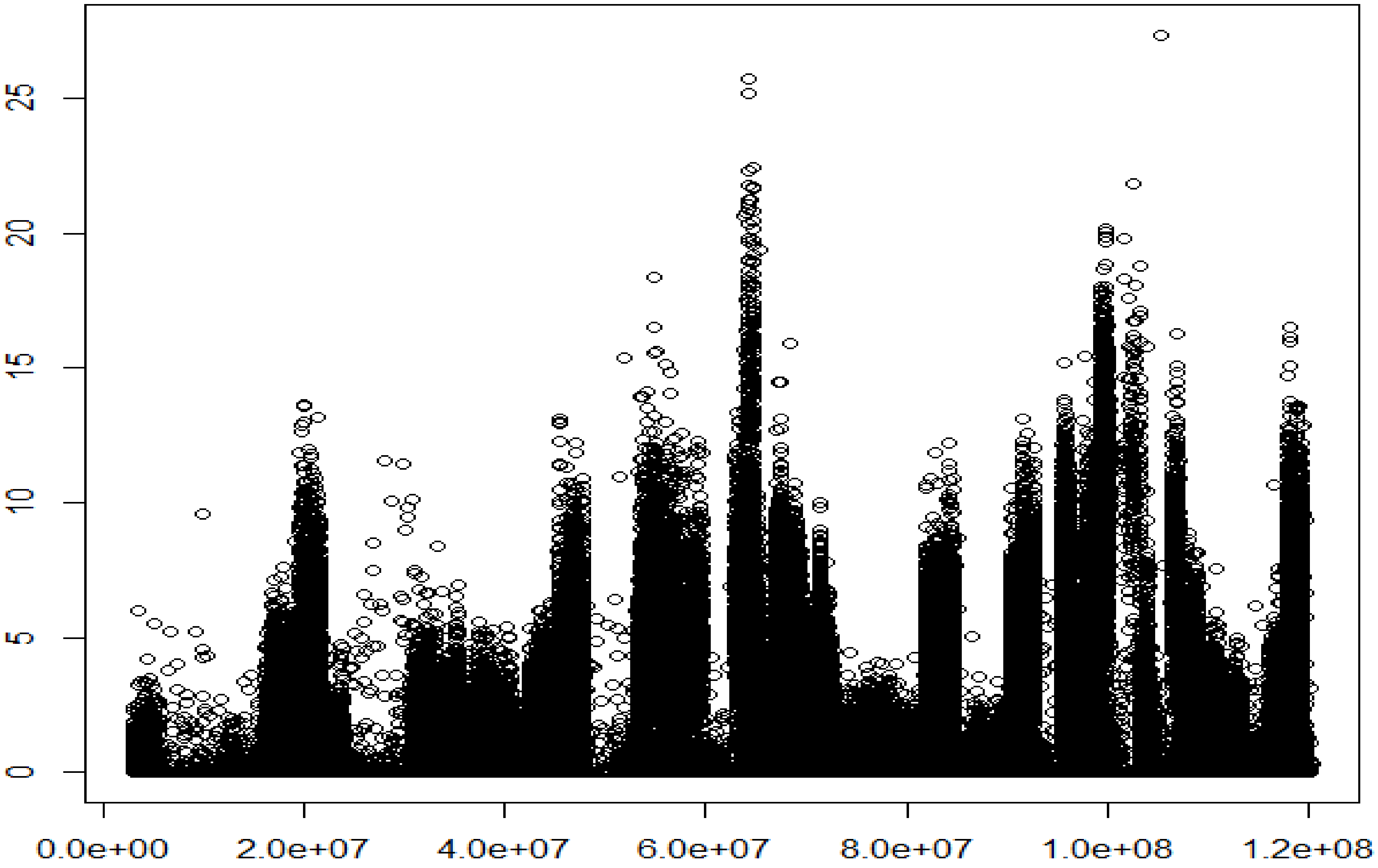


**Figure S13**. Manhattan plot of the results from fisher's exact test for difference in allele frequency of 187,440 variants on Mouse chromosome 13. Horizontal axis, location (base pair); Vertical axis, -ln(p) value. Threshold value for significance = -ln(0.05/4,147,085) = 18.23.

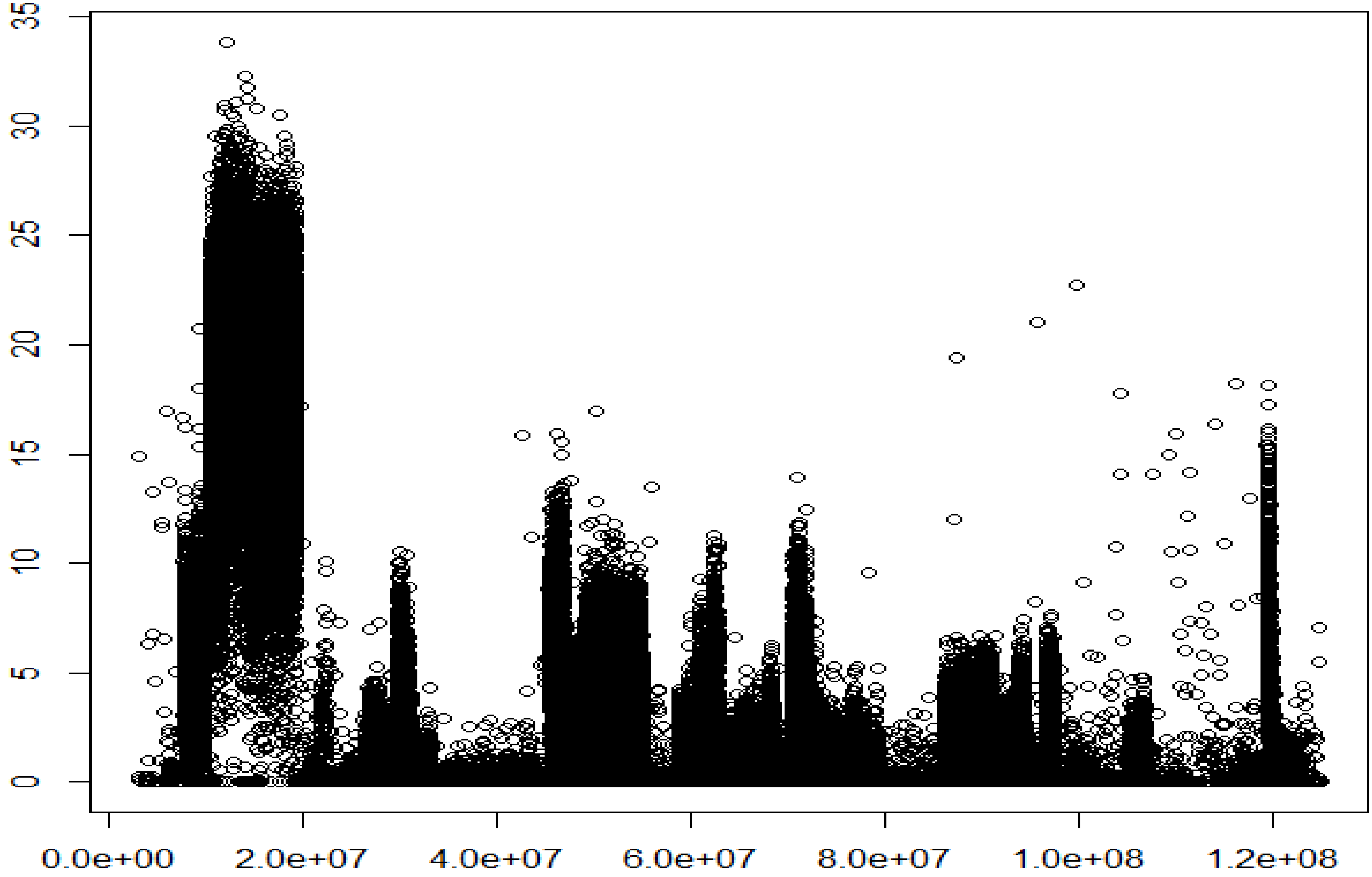


**Figure S14**. Manhattan plot of the results from fisher's exact test for difference in allele frequency of 235,392 variants on Mouse chromosome 14. Horizontal axis, location (base pair); Vertical axis, -ln(p) value. Threshold value for significance = -ln(0.05/4,147,085) = 18.23.

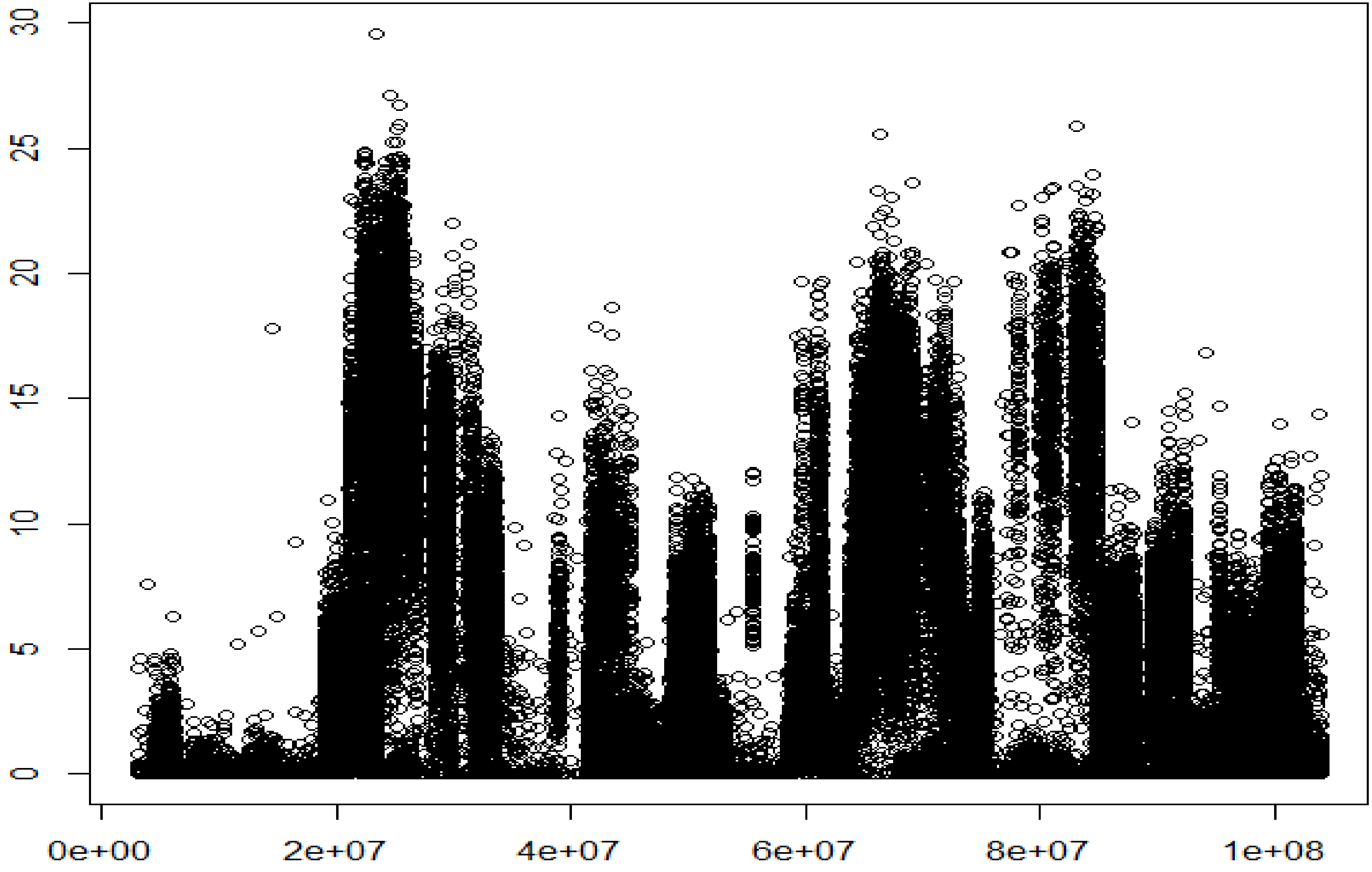


**Figure S15**. Manhattan plot of the results from fisher's exact test for difference in allele frequency of 144,143 variants on Mouse chromosome 15. Horizontal axis, location (base pair); Vertical axis, -ln(p) value. Threshold value for significance = -ln(0.05/4,147,085) = 18.23.

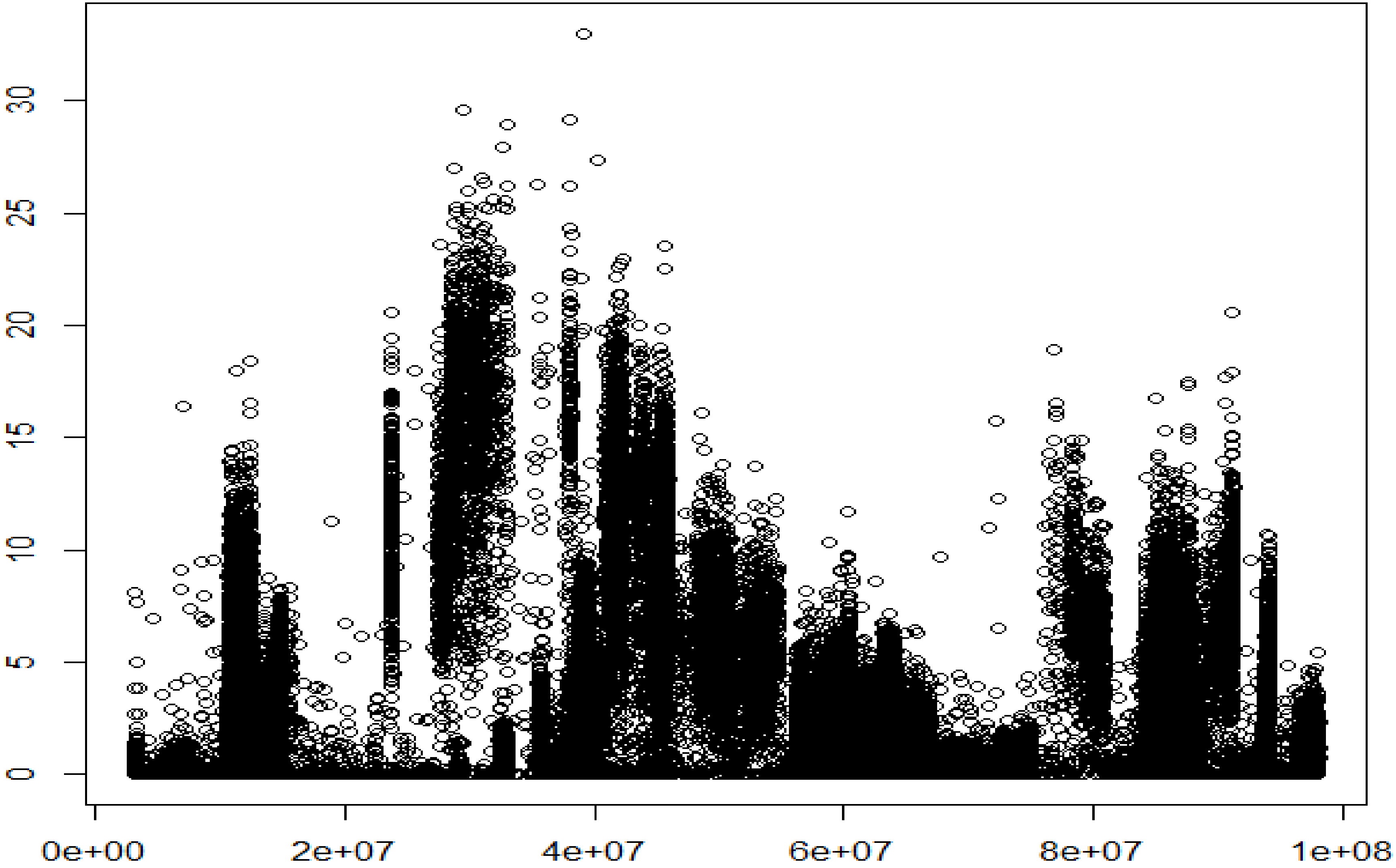


**Figure S16**. Manhattan plot of the results from fisher's exact test for difference in allele frequency of 96,525 variants on Mouse chromosome 16. Horizontal axis, location (base pair); Vertical axis, -ln(p) value. Threshold value for significance = -ln(0.05/4,147,085) = 18.23.

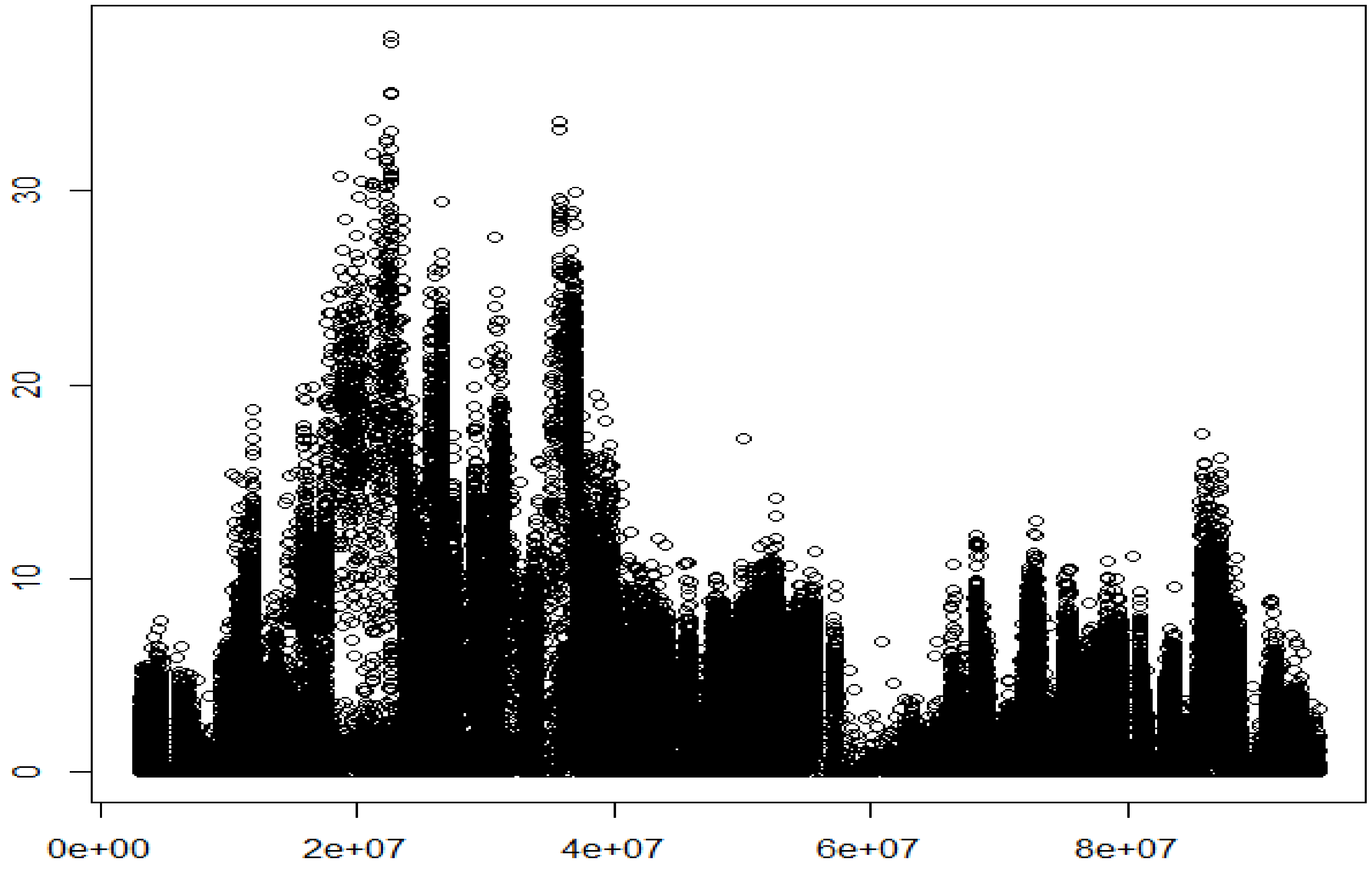


**Figure S17**. Manhattan plot of the results from fisher's exact test for difference in allele frequency of 163,233 variants on Mouse chromosome 17. Horizontal axis, location (base pair); Vertical axis, -ln(p) value. Threshold value for significance = -ln(0.05/4,147,085) = 18.23.

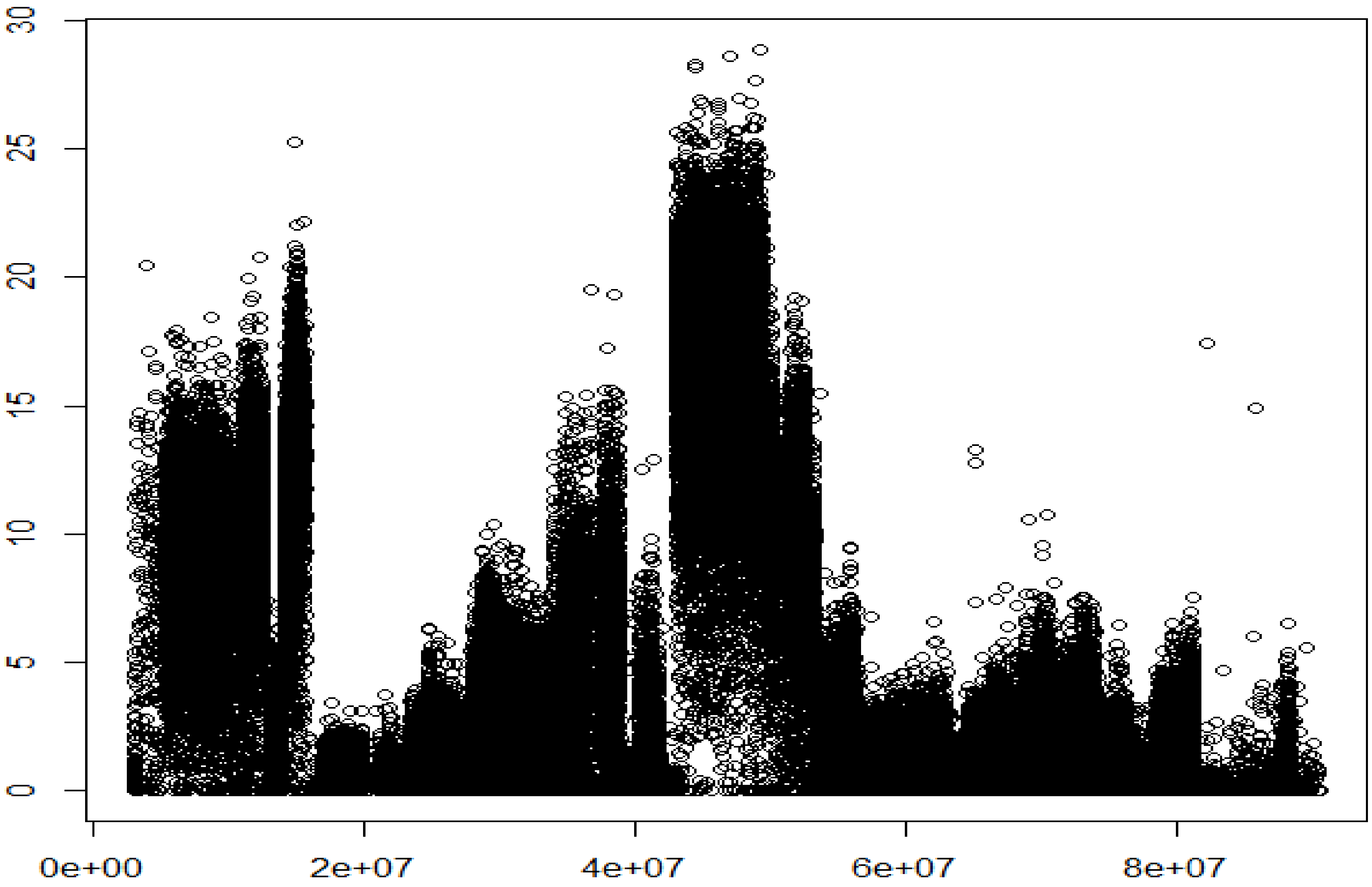


**Figure S18**. Manhattan plot of the results from fisher's exact test for difference in allele frequency of 176,065 variants on Mouse chromosome 18. Horizontal axis, location (base pair); Vertical axis, -ln(p) value. Threshold value for significance = -ln(0.05/4,147,085) = 18.23.

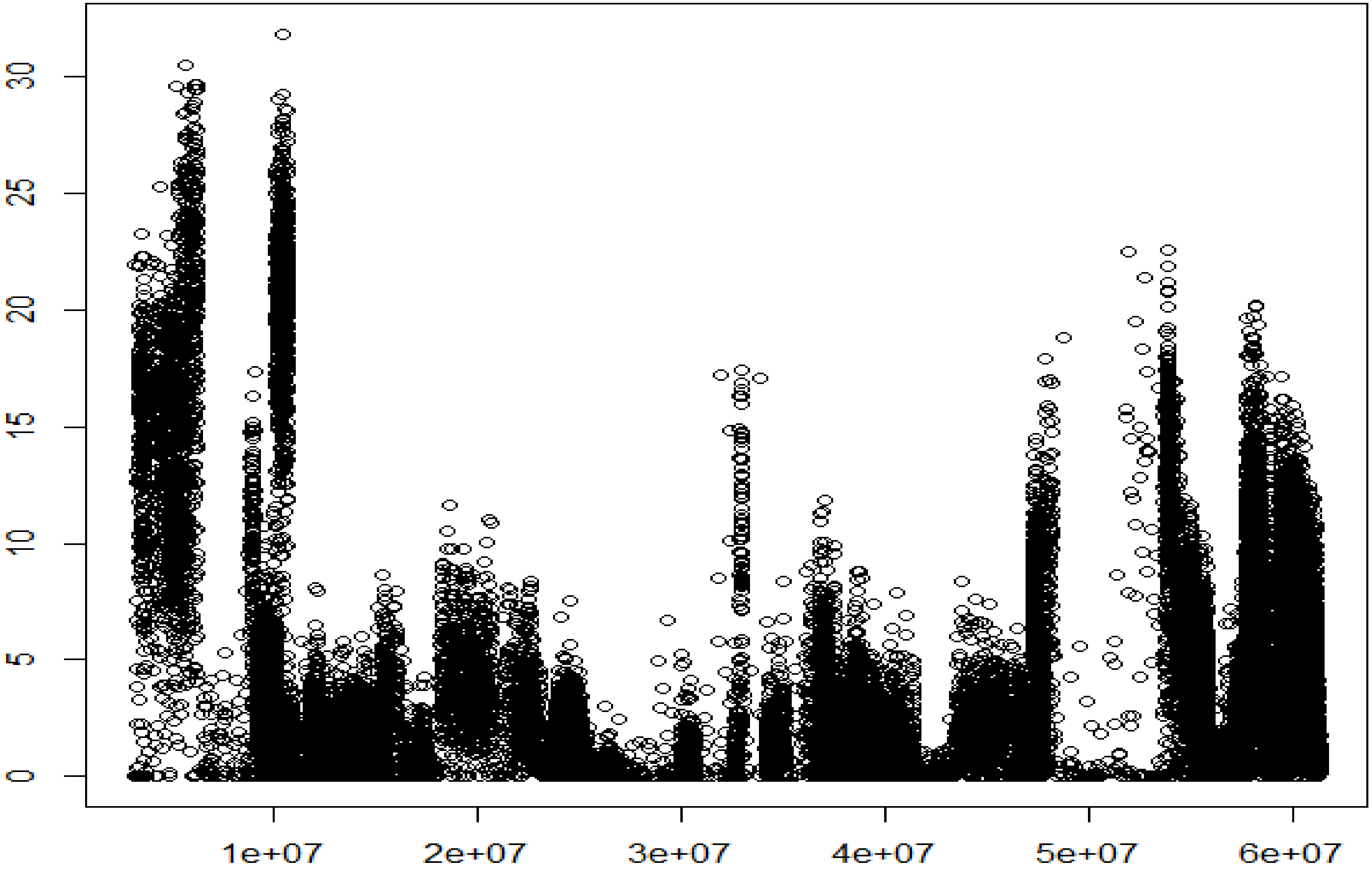


**Figure S19**. Manhattan plot of the results from fisher’s exact test for difference in allele frequency of 45,914 variants on Mouse chromosome 19. Horizontal axis, location (base pair); Vertical axis, -ln(p) value. Threshold value for significance = -ln(0.05/4,147,085) = 18.23.
